# A Liquid-to-Solid Phase Transition Enhances the Catalytic Activity of SARM1

**DOI:** 10.1101/2020.08.28.272377

**Authors:** Heather S. Loring, Paul R. Thompson

## Abstract

Sterile alpha and toll/interleukin receptor (TIR) motif–containing protein 1 (SARM1) is a neuronally expressed NAD^+^ glycohydrolase whose activity is increased in response to various stressors. The consequent depletion of NAD^+^ triggers axonal degeneration (i.e., Wallerian degeneration), which is a characteristic feature of neurological diseases, including peripheral neuropathies and traumatic brain injury. Notably, SARM1 knockout mice show minimal degeneration in models of peripheral neuropathy and traumatic brain injury, making SARM1 a promising therapeutic target. However, the development of SARM1 inhibitors has been challenging as the purified enzyme is largely inactive. Herein, we report that SARM1 activity is increased ∼2000–fold by a liquid-to-solid phase transition. These findings provide critical insights into SARM1 biochemistry with important implications for the situation *in vivo*. Moreover, they will facilitate the discovery of novel SARM1–targeted therapeutics.

**Graphical Abstract:** 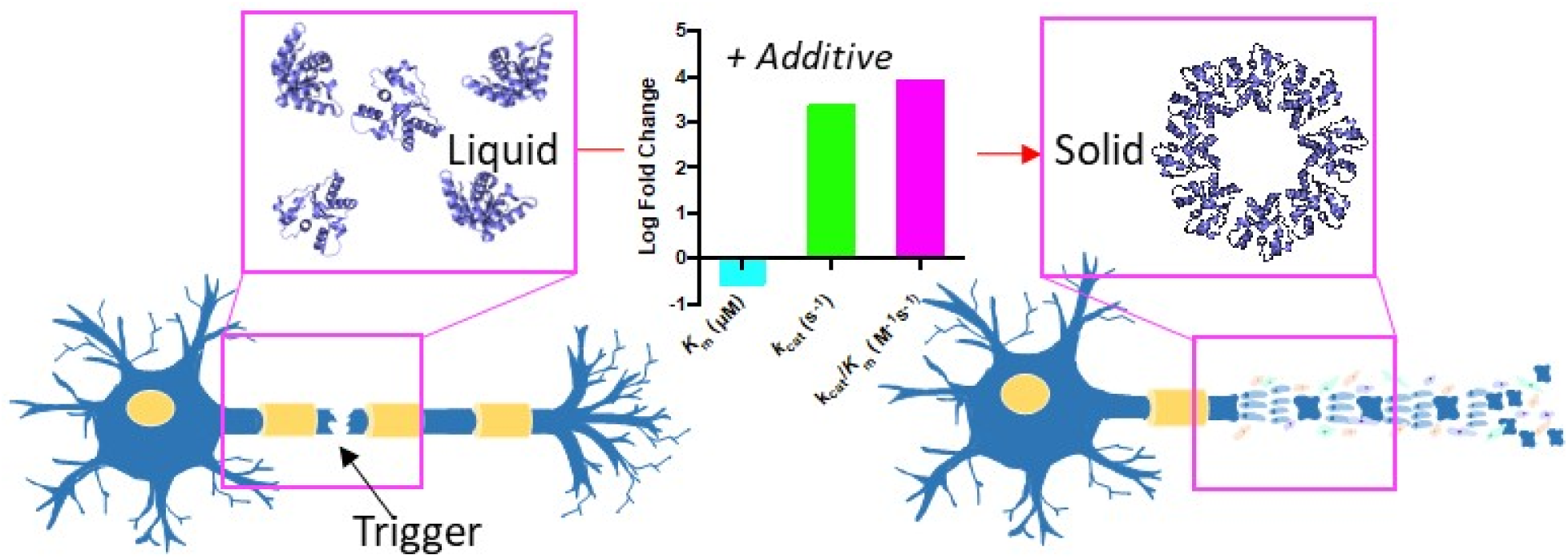

---

Despite diverse clinical manifestations, axonal degeneration is an underlying feature of traumatic brain injury, peripheral neuropathies, and other neurodegenerative diseases, including Alzheimer’s disease, Huntington’s disease, and Parkinson’s disease. These diseases account for extensive morbidity and mortality as there are no approved therapies. SARM1 has recently emerged as a promising therapeutic target for diseases associated with axonal degeneration because SARM1 knockdown or knockout prevents axonal degeneration and disease pathology (Geisler et al., 2016; Henninger et al., 2016; Loring and Thompson, 2020a).

SARM1 was originally found to play a role in axonal degeneration when it was identified in a screen that selected for mutants that suppress injury–induced axonal degeneration (Osterloh *et al*., 2012). However, it was unknown how the loss of SARM1 prevented neurodegeneration post–injury. Subsequently, SARM1 was shown to possess enzymatic activity and catalyze the hydrolysis of NAD^+^ (Figure 1A) (Essuman et al., 2017). Depletion of the NAD^+^ pool promotes degeneration whereas inactive variants of SARM1 or NAD^+^ supplementation prevent degeneration after injury (Essuman et al., 2017; Gerdts et al., 2015; Wang et al., 2015).

**Figure 1.**
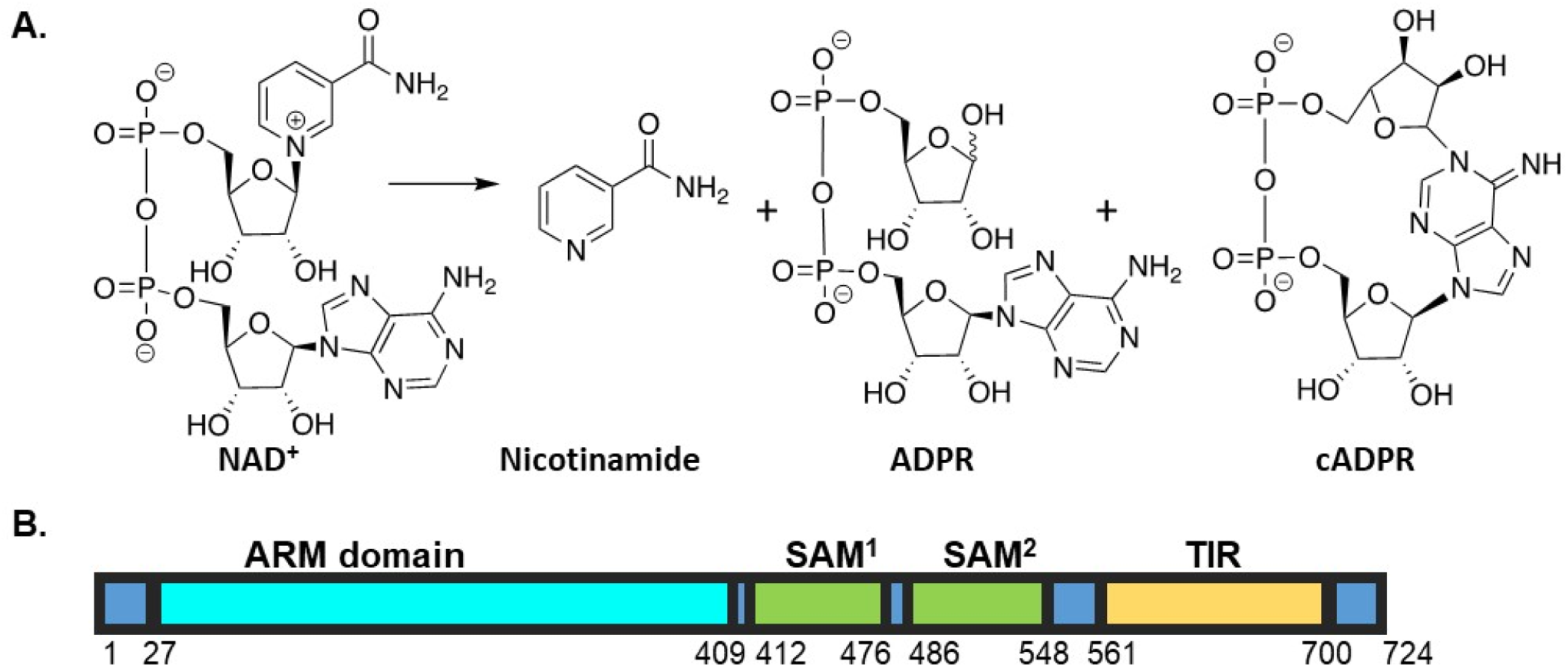
SARM1 Reaction and Domain Structure. (A) SARM1-mediated hydrolysis of NAD^+^ results in the production of nicotinamide and a mixture of ADPR and cADPR. (B) Domain architecture of SARM1.

SARM1 consists of three domains: a HEAT/armadillo (ARM) domain, two tandem sterile alpha motif (SAM) domains, and a C-terminal toll interleukin receptor (TIR) domain that catalyzes NAD^+^ hydrolysis (Figure 1B). The TIR domain contains a BB loop (Residues 594–605), which is thought to promote self– association, and a SARM1–specific (SS) loop (Residues 622–635), which is a characteristic feature of catalytically active TIR domains; TIR domains lacking this loop do not possess enzymatic activity (Carlsson et al., 2016; Horsefield et al., 2019; Loring and Thompson, 2020a; Summers et al., 2016). Additionally, E642 (within the TIR domain and adjacent to the SS loop) is thought to be a key catalytic residue based on structural alignments with other nucleotide hydrolases (Essuman et al., 2017; Loring and Thompson, 2020a). Under healthy conditions SARM1 activity needs to be inhibited. One widely accepted model involves the ARM domain inhibiting catalysis via an intramolecular interaction with the catalytic TIR domain. In response to injury or disease, the enzyme is thought to undergo a conformational change that relieves this interaction, which allows the SAM domains to multimerize and activate NAD^+^ hydrolysis (Gerdts et al., 2013; Summers et al., 2016). This model suggests that targeting multimerization or NAD^+^ hydrolysis may be viable approaches for inhibitor development.

Although SARM1 inhibitors hold therapeutic promise, efforts to study the purified enzyme have been complicated by the fact that SARM1 appears to lose activity during purification and only shows measurable NAD^+^ hydrolase activity when assayed at a final concentration of ≥ 20 µM (Horsefield et al., 2019; Loring et al., 2020a). By contrast, enzymatic activity is 800–fold higher in lysates or when activity is measured on–bead (Essuman et al., 2017; Horsefield et al., 2019; Loring et al., 2020a). These data indicate that the pure protein does not fully recapitulate all features of SARM1 catalysis. Herein we report that a liquid-to-solid phase transition increases SARM1 activity by ∼2000–fold to levels approaching that observed in lysates. By exploiting this finding, we report the first detailed studies of SARM1 catalysis.

## Results and Discussion

We previously established that SARM1 loses activity during purification, and is essentially inactive unless purified to high micromolar concentrations (Figure 2A) (Loring et al., 2020a). Initially, we hypothesized that the loss in activity could be explained by the loss of an essential element during enzyme purification. However, the addition of purified SARM1 to cell lysates did not activate the enzyme (Figure 2B). We then hypothesized that pure SARM1 adopts an inactive conformation. To investigate this hypothesis, we evaluated the effect of enzyme concentration (5–35 µM) on the rates of catalysis. Notably, the concentration dependence has parabolic character, increasing with upward curvature, as opposed to linearly, suggesting the formation of a more active oligomer (Figure 2C). The fact that pure SARM1 regains activity upon concentration, proves that an essential cofactor is not lost during purification. To investigate the oligomerization status of the SARM1 TIR domain, we next performed analytical ultracentrifugation (AUC). SARM1 was largely monomeric (∼90%), as indicated by an S–value of 1.9–2.0S (Figure 2D). A smaller peak at ∼3.9–4.2S, which showed rightward tailing indicates that the remaining ∼10% exists as a trimer or tetramer (∼71–88 kDa) with a smaller fraction in higher ordered forms. Similar oligomerization levels were observed at the two different SARM1 concentrations tested (Figure 2D).

**Figure 2.**
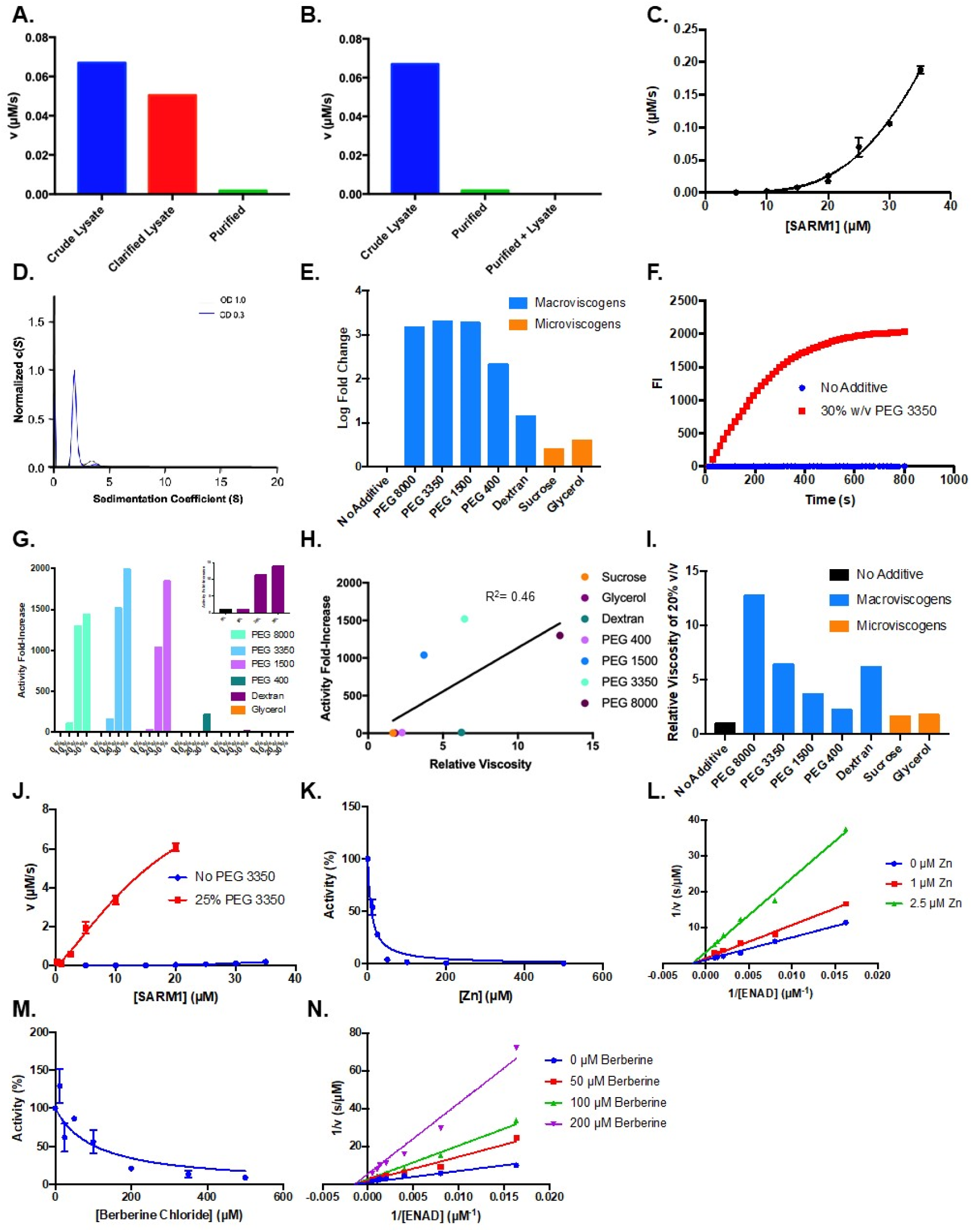
Effect of Crowding Agents on Activity. (A) Activity of crude lysate, clarified lysate, and elution during SARM1 purification. (B) Activity of crude lysates, pure protein and pure protein added back to C43 lysates. (C) Dose dependence of SARM1 TIR domain activity from 5– 35 µM with 1 mM ENAD. (D) AUC of SARM1 at an OD_280_ of 0.3 and 1.0. (E) Log fold change in SARM1 (10 µM) activity in buffer compared to 30% of crowding agent and 1 mM ENAD. (F) Representative time course of SARM1 TIR domain with and without 30% w/v PEG 3350 at 2 mM ENAD and 20 µM SARM1. (G) Dose dependence of SARM1 TIR domain activity from 0 – 30 % crowding agent and 1 mM ENAD. (H) Activity of SARM1 (10 µM) at different viscosities of 20% w/v solutions. (I) Relative viscosities at 20% w/v of different solutions. (J) Dose dependence of SARM1 TIR domain activity from 300 nM to 35 µM with and without PEG. (K) Zinc inhibition of SARM1 (2.5 µM) plus 25% PEG 3350 with 100 µM ENAD. (L) Mechanism of zinc inhibition of SARM1 (2.5 µM) in the presence of 25% PEG 3350. (M) Berberine chloride inhibition of SARM1 (2.5 µM) plus 25% PEG 3350 with 100 µM ENAD. (N) Mechanism of berberine chloride inhibition of SARM1 (2.5 µM) in the presence of 25% PEG 3350.

Having shown that SARM1 activity increases nonlinearly with concentration, we next hypothesized that molecular crowding might activate the enzyme. To test this hypothesis, we measured the activity of SARM1 in the presence of several macroviscogens (i.e., PEG 8000, 3350, 1500, 400, and Dextran) and microviscogens (i.e., Sucrose and Glycerol). Microviscogens are often used as protein stabilizers and affect the ability of small molecules to diffuse in solution, whereas macroviscogens alter the free volume available for a molecule to move within a solution with little effect on diffusion (Blacklow et al., 1988; Gadda and Sobrado, 2018; Hagen, 2010). Therefore, microviscogens can reveal effects on diffusional processes, whereas macroviscogens affect molecular crowding (Gadda and Sobrado, 2018). Notably, we found that all the macroviscogens tested increased activity while the microviscogens had negligible effects (Figure 2E). Specifically, PEGs 8000, 3350, 1500, and 400 increased activity by 1500, 2000, 1900, and 200–fold, respectively. By contrast, dextran (36–50 kDa) only increased activity by 14–fold (Figure 2E). To better visualize the effect of PEG 3350 on the rate of the reaction, representative progress curves are shown at 20 µM SARM1 with and without PEG (Figure 2F). The 20 µM reaction with PEG 3350 plateaus after ∼ 350 s, whereas the fluorescence of the reaction without PEG increases at a much slower rate. In total, these data indicate that a subset of macroviscogens dramatically increase SARM1 activity by up to 2000–fold.

To determine how macroviscogens increase SARM1 catalysis, the activity of SARM1 (10 µM final) was next measured at 0, 10, 20, and 30% w/v of the macroviscogens and glycerol. Consistent with our hypothesis, we observed a dose–dependent increase in activity for all the macroviscogens tested (Figure 2G). Notably, activity increases according to macroviscogen size until a critical point is reached where PEG 1500 and PEG 3350 have similar effects (Figure 2G). Note that the effect on activity is not purely due to a change in viscosity, as the enhancement in activity shows a poor correlation (R^2^= 0.46) with viscosity (Figure 2H). For example, when dextran and PEG 3350 are assayed as similar relative viscosities, the fold activation is dramatically different (Figure 2H–I). Additionally, when the viscosity of glycerol was compared to that of PEG 3350, we found that glycerol did not enhance activity to the same degree as PEG 3350, despite having a similar relative viscosity (Figure S1A–B).

As PEG 3350 showed the greatest ability to enhance activity, we proceeded to investigate its effect on SARM1 activity further. Initially, we evaluated the effect of enzyme concentration on activity in the presence and absence of PEG 3350 (Figure 2J). SARM1 concentration was varied from 300 nM–20 µM with 25% w/v PEG 3350 and from 5–35 µM in the absence of PEG 3350. Our data shows that the addition of 25% w/v PEG 3350 accelerates the rate of reaction 7,700–fold at 5 µM, 1,400–fold at 10 µM, and ∼280–fold at 20 µM of SARM1 (Figure 2J). The fact that the increase in activity changes with enzyme concentration, indicates that the effect of PEG 3350 is dependent on enzyme concentration. Furthermore, SARM1 activity appears to increase almost linearly with respect to concentration in the presence of PEG after a slight lag phase, but parabolically in its absence (Figure 2J). This trend suggests that in the absence of PEG, there is a gradual build up in activity as the concentration of SARM1 increases, until the concentration surpasses a critical threshold allowing for increased transient interactions and a consequent large increase in activity (Figure 2J). By contrast, the addition of PEG reduces the concentration of enzyme needed for activity. One potential explanation is that PEG induces molecular crowding by excluding water from the protein surfaces, thereby promoting multimerization and catalysis. Along these lines, we note that in the absence of PEG the Hill coefficient is 3.8, indicating a high degree of cooperativity. Whereas, in the presence of PEG (0–25%) the Hill coefficient decreases to 1.3, suggesting that PEG decreases the dependence on the cooperative formation of an active multimer (Figure S1C–D).

To assess whether the natural behavior of SARM1 was maintained in the presence of PEG 3350, we evaluated the potency of several metals that were previously shown to inhibit SARM1 when assayed in lysates (Loring et al., 2020a; Loring and Thompson, 2020b). Notably, NiCl_2_ and ZnCl_2_ completely eliminated activity in the presence of PEG (Figure S1E, (Loring et al., 2020a)). This result is consistent with our previous data (Loring et al., 2020a). Expanding on these studies, we also evaluated several other metal ions including CaCl_2_, CdCl_2_, CoCl_2_, CuCl_2_, MgCl_2_, and MnCl_2_ (Figure S1E). Of note, CdCl_2_, CoCl_2_, and CuCl_2_, decreased activity in the presence of the pure protein but not in lysates. This inconsistency could be due to elements in the lysate preferentially sequestering these metals and thereby preventing their ability to act on SARM1. Given the high potency of CdCl_2_ and CuCl_2_, we used these inhibitors to titrate the enzyme and determine the percentage of active enzyme (Figure S1F). Based on this data, all the enzyme appears to be active. Although speculative, the fact that thiophilic metals (i.e., CdCl_2_, CoCl_2_, CuCl_2_, NiCl_2_, and ZnCl_2_) inhibit SARM1 suggests the presence of a catalytically important thiol. Along these lines, we previously showed that C629 and C635, which are present in the SARM1 specific loop, are critical for activity in the absence of PEG (Loring and Thompson, 2020b).

We next determined an IC_50_ value for ZnCl_2_ as this compound was previously shown to potently inhibit the SARM1 TIR domain. Consistent with our prior data, the IC_50_ value obtained in the presence of PEG is 10 ± 1 µM, which is very similar to that in lysates (10 ± 3 µM) (Loring et al., 2020a), and with purified protein without PEG (10 ± 1 µM) (Loring and Thompson, 2020b) (Figure 2K, Table 1). To further confirm that the addition of PEG 3350 was not introducing any confounding variables, we repeated the mechanism of inhibition studies with ZnCl_2_. As before, we found that ZnCl_2_ inhibits SARM1 noncompetitively with *K*_i_ value of 1.0 ± 0.1 µM. This value is similar to the value obtained when lysate derived SARM1 was assayed (*K*_i_ = 3.3 ± 0.1 µM (Figure 2L) (Loring and Thompson, 2020b)).

**Table 1.** Comparison of Inhibitor Potency.

| | Lysate $IC_{50}$<br>($\mu M$)<sup>a</sup> | Pure Protein $IC_{50}$<br>($\mu M$)<sup>b</sup> | Pure Protein + PEG $IC_{50}$<br>($\mu M$)<sup>c</sup> |
| --- | --- | --- | --- |
| Zinc | $10 \pm 3$ | $10 \pm 1$ | $10 \pm 1$ |
| Berberine Chloride | $140 \pm 20$ | $140 \pm 20$ | $140 \pm 30$ |

Assays were performed at SARM1 concentrations of 300 nM<sup>a</sup>, 20 $\mu M$ <sup>b</sup>, and 2.5 $\mu M$ <sup>c</sup>.

To additionally confirm that PEG does not impact the natural behavior of SARM1, we tested berberine chloride, an inhibitor previously identified from a high throughput screen (Loring and Thompson, 2020b). Berberine chloride inhibited SARM1 in the presence of PEG with an IC_50_ value of 140 ± 30 µM. This value is quite similar to its potency in lysates (140 ± 20 µM) and when assayed with purified SARM1 without PEG 3350 (140 ± 20 µM) (Figure 2M and Table 1) (Loring and Thompson, 2020b). As before, we found that berberine chloride also inhibits SARM1 noncompetitively in the presence of PEG 3350 with a *K*_i_ value of 120 ± 10 µM. This value and inhibition pattern are identical to that obtained when SARM1 was assayed in lysates (*K*_i_ = 120 ± 10 µM, Figure 2N (Loring and Thompson, 2020b)).

We next sought to investigate how PEG enhances the activity of SARM1. First, we performed AUC in the presence of PEG to investigate whether PEG was inducing the formation of a higher ordered oligomer. However, we did not detect any protein in the sample. Since PEGs are typically used as precipitants, we hypothesized that our inability to detect SARM1 was due to the precipitation of SARM1 in the presence of PEG 3350. Consistent with this hypothesis, we found that the addition of PEG 3350 (25% w/v) results in the formation of an active precipitate, whereas the protein in buffer without PEG remains soluble (Figure 3A). These data indicate that the observed rate enhancement is driven by a liquid-to-solid phase transition. Next, we evaluated the effect of protein and PEG concentration on this liquid-to-solid phase transition. We found that both enzyme (5–20 µM) and PEG (0–25%) concentration enhanced precipitate formation (Figure 3B–C). As the concentration of SARM1 in the supernatant decreases, the precipitate concentration increases with a corresponding increase in enzyme activity. This effect is evident by the absence of a pellet for 0% PEG, a faint band at 10% PEG, and darker bands at 17.5% and 25% PEG at 5, 10, and 20 µM SARM1, and the inverse trend for the supernatant (Figure 3C). To determine whether other precipitants also increase SARM1 activity, we screened a panel of common precipitants and identified citrate as another highly potent activator of SARM1 activity (Figure S1G).

**Figure 3.**
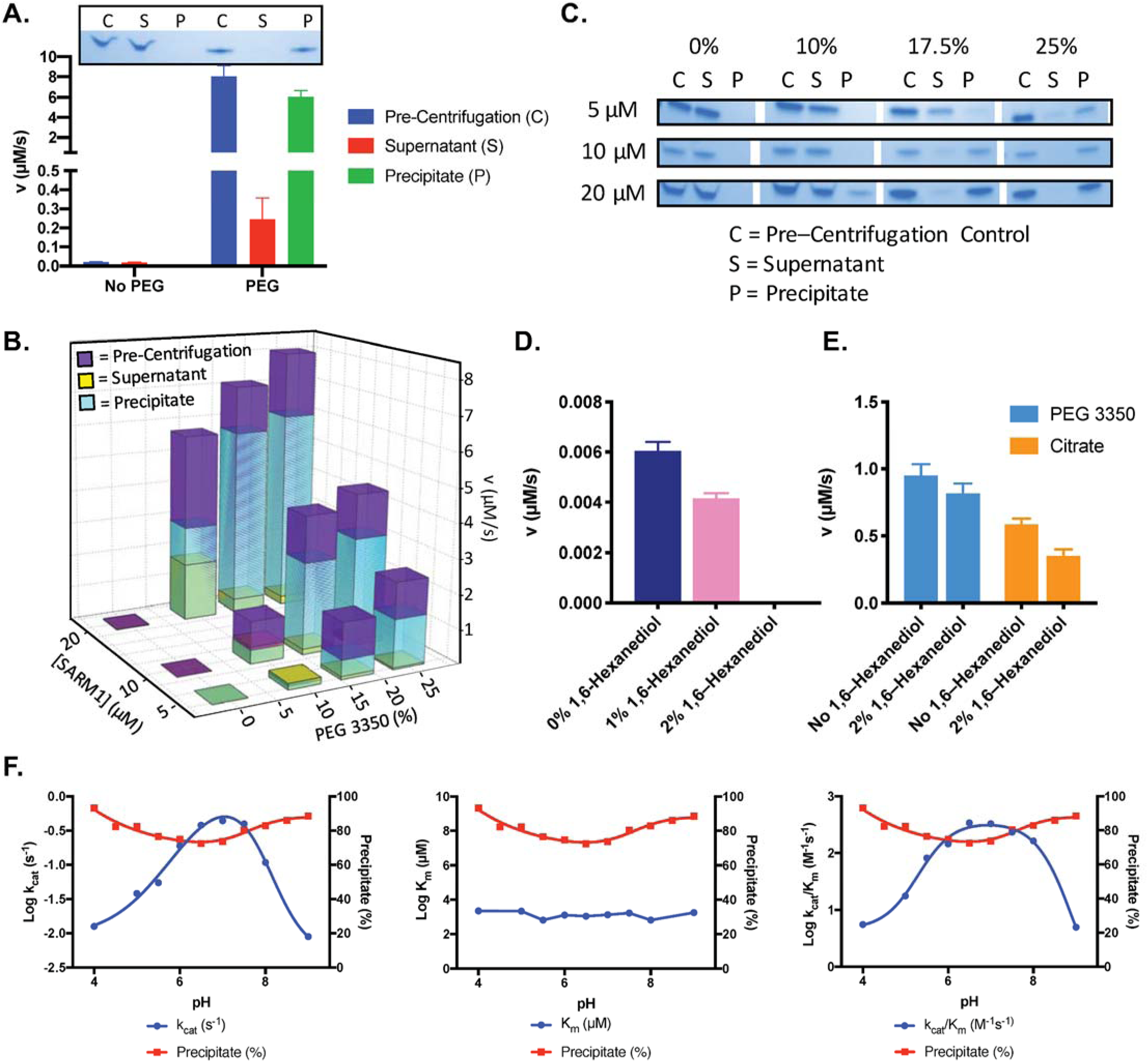
PEG 3350 Induces a Liquid to Solid Phase Transition. (A) Activity of pre-centrifugation, supernatant, and precipitate fractions obtained before and after treatment of 20 µM SARM1 with and without 25% PEG 3350. Coomassie gels demonstrate the presence of SARM1 in each fraction. (B) Activity of pre-centrifugation, supernatant, and precipitate fractions obtained after treating SARM1 (5 µM, 10 µM, and 20 µM) with 0%, 10%, 17.5%, and 25% PEG 3350. (C) Coomassie stained gels demonstrate the presence of SARM1 in each fraction analyzed in panel B. (D) Effect of 1,6–Hexanediol on pure SARM1 (20 µM) activity. (E) Effect of 2% 1,6–hexanediol treatment on SARM1 activity in the presence of 25% PEG or 500 mM sodium citrate. (F) Effect of pH on the steady state kinetics and precipitate formation. Data is from an average of three experiments each conducted in duplicate. Full gels provided in Figure S3A-D.

To explore the nature of the interactions involved in precipitate formation, we tested the effect of 1,6– hexanediol, an aliphatic alcohol that is often used to distinguish between hydrophobic interactions involved in liquid-to-liquid phase separations and liquid-to-solid phase transitions (Kroschwald et al., 2015; Patel et al., 2007; Peskett et al., 2018). We found that while 1,6–hexanediol inhibited the activity of pure protein alone, it did not substantially affect the activity of SARM1 in the presence of PEG or sodium citrate (Figure 3D–E). This data suggests that the activity of the pure protein is dominated by weak hydrophobic interactions or a liquid-to-liquid phase separation (Figure 3D). On the other hand, the fact that 1,6–hexanediol does not eliminate the activity of pure protein in the presence of PEG or sodium citrate, supports SARM1 undergoing a liquid-to-solid phase transition in these cases (Figure 3E). To confirm that precipitate formation is reversible, we performed the centrifugation experiment in the presence of PEG or sodium citrate and tested the activity of the fractions (pre-centrifugation, supernatant, precipitate resuspended in buffer, and precipitate resuspended with buffer plus additive) (Figure S1H). The lack of both a precipitate and activity in the precipitate resuspended in buffer confirm that precipitation is reversible.

Having established that both PEG 3350 and sodium citrate enhance SARM1–mediated NAD^+^ hydrolysis via a liquid-to-solid phase transition, we sought to exploit this property to perform the first detailed analysis of SARM1 catalysis. We first determined how pH affects activity and precipitate formation. We found that the catalytic efficiency (*k*_cat_/*K*_m_) peaks from pH 6–8, which is primarily due to an effect on *k*_cat_, as the *K*_m_ value remains nearly constant across the ranges tested (Figure 3F). Furthermore, the effect of pH on activity appears to be independent of precipitation as the precipitate percentage varies by only 10% across the range tested. Notably, the pH profile for *k*_cat_ is bell-shaped with p*K*_a_ values of 4.9 ± 0.2 and 8.8 ± 0.2 for the ascending and descending limbs, respectively. The p*K*_a_ value for the ascending limb is consistent with the previous suggestion that a glutamate, likely E642, acts as a key catalytic residue (Essuman et al., 2017).

We next evaluated the effect of PEG on the steady state kinetic parameters in the presence of 10 µM SARM1. In the presence of 25% PEG, the *K*_m_ decreases 3.3–fold, whereas *k*_cat_ and *k*_cat_/*K*_m_ increase by 2000– and 8000–fold, respectively (Figure 4A). The fact that the increase in *k*_cat_/*K*_m_ is dominated by an increase in *k*_cat_, suggests that PEG promotes multimerization and activity via precipitate formation. To further interrogate how PEG affects the kinetic activity of SARM1, we performed detailed steady state kinetic experiments at various PEG concentrations from 0–25% (Figure 4B). We found that increasing concentrations of PEG 3350 cause the *K*_m_ value to decrease and level off at ∼500 µM (Figure 4C) and the *k*_cat_ to increase almost linearly (Figure 4D). The catalytic efficiency also increases and plateaus at ∼1700 M^-1^s^-1^ (Figure 4E, Table S1). Interestingly, this value is nearly identical to that recorded in the lysate and on–bead (1500 M^−1^s^−1^) (Essuman et al., 2017; Loring et al., 2020a) (Figure 4B–E, Table 2).

**Table 2.** Comparison of Steady-State Kinetic Parameters for the SARM1 TIR domain.

| | 2.5 μM<br>PEG | Lysate<sup>a</sup> | On-Bead<sup>a</sup> | Literature<sup>b</sup> |
| --- | --- | --- | --- | --- |
| $K_m$ (μM) | $230 \pm 30$ | $40 \pm 10$ | $28 \pm 6$ | 24 |
| $k_{cat}$ ($s^{-1}$) | $0.51 \pm 0.01$ | $0.06 \pm 0.03$ | $0.04 \pm 0.03$ | 0.17 |
| $k_{cat}/K_m$ ($M^{-1}s^{-1}$) | $2200 \pm 30$ | $1500 \pm 300$ | 1500 | 7083 |

<sup>a</sup>(Loring et al., 2020a)<sup>b</sup>Reported by Essuman *et al.* 2017 with SARM1 on-bead by LC-MS.

**Figure 4.**
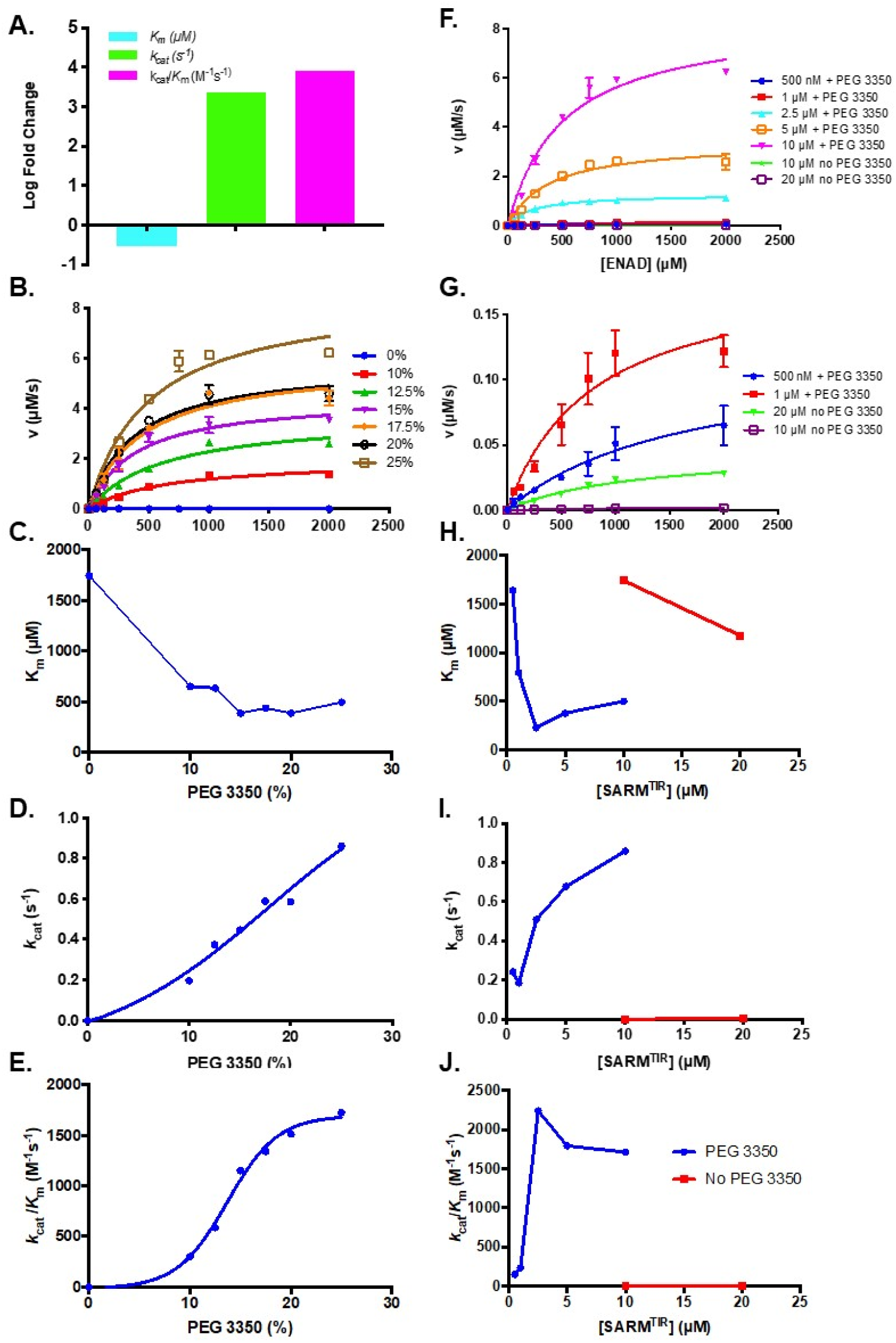
Effect of PEG 3350 Concentration on the Steady State Kinetics of ENAD Hydrolysis. (A) Steady state kinetic analysis at 10 µM SARM1 with and without addition of PEG 3350 (25%). (B) Steady state kinetic analysis of SARM1 (10 µM) with PEG 3350 (0–25% w/v) (C) *K*_m_, (D) *k*_cat_, and (E) *k*_cat_/*K*_m_ values at each concentration. (F) Steady state kinetic analysis of SARM1 from 500 nM – 10 µM with and without PEG 3350 at 10 and 20 µM from 0 – 2 mM ENAD fit to the Michaelis Menten equation. (G) Blow up of steady state kinetic analysis of SARM1 at 500 nM and 1 µM with PEG 3350, and 10 and 20 µM without PEG 3350 with ENAD from 0-2 mM. Fitting to Michaelis Menten equation gives (H) *K*_m_, (I) *k*_cat_, and (J) *k*_cat_/*K*_m_ values at each concentration.

We also performed steady state kinetic analyses at different concentrations of SARM1 in the absence and presence of 25% w/v PEG 3350 (Figure 4F–J). As with increasing PEG concentration, an increase in SARM1 concentration led to a small decrease in *K*_m_ and an increase in *k*_cat_ (Figure 4F–J, Table S2), such that the *k*_cat_/*K*_m_ values are comparable to those obtained for the enzyme assayed in lysate (Loring et al., 2020a). Of particular note is the fact that PEG makes kinetic studies feasible at 500 nM of SARM1, which is near the concentration studied in lysates (300 nM) (Loring et al., 2020a). At 500 nM of SARM1 with PEG, the *K*_m_ is 1600 µM, the *k*_cat_ is 0.24 s^−1^, and the catalytic efficiency is 150 M^−1^s^−1^ (Figure 4F–J, Table S2). Notably, the *K*_m_ value at 500 nM with PEG is similar to that at 20 µM SARM1 without PEG (1200 µM), however the *k*_cat_ is ∼100–fold greater in the presence of PEG (Figure 4F–J, Table S2). These data suggest that by inducing a liquid-to-solid phase transition and forming an ordered oligomer, PEG drastically enhances enzymatic turnover, but has little effect on substrate binding. Although the *K*_m_ value slightly decreases as SARM1 or PEG concentration increases, it remains much higher than the lysate value of 40 ± 10 µM (Loring et al., 2020a), indicating that PEG/SARM1 concentration does not completely recapitulate SARM1 activity in the lysate or on–bead (Figure 4, Table 2). The similarities in *k*_cat_/*K*_m_, however, suggest compensation (i.e., alterations in rate limiting steps) between these two parameters. Moreover, the fact that *k*_cat_ is directly correlated with enzyme and PEG concentration, suggests that turnover rate is largely dependent on the multimeric state and the formation of an ordered oligomer.

Next, we evaluated the product specificity of SARM1 with PEG and sodium citrate. As with pure enzyme, we found that the addition of PEG or citrate, led to a decrease in NAD^+^ levels with time and increased levels of nicotinamide as well as a mixture of ADPR and cADPR (Figure 5A–B, S2A–B). Of note, the ADPR to cADPR ratios are consistent over time under each condition (Figure 5C), however, the addition of PEG increases the production of ADPR relative to cADPR from a ratio of 10 ± 2 to a ratio of 16 ± 1. This ratio is further increased in the presence of citrate to 23 ± 3 (Figure 5D). We hypothesize that this could be due to the multimeric state of the enzyme constraining NAD^+^ in such a position that it favors hydrolysis by water as compared to reaction with the N1 of the adenosine, which is required for cyclization. We also evaluated whether SARM1 could cleave cADPR in the presence or absence of additive (Figure S2C). Consistent with our prior finding that SARM1 follows an ordered uni–bi kinetic mechanism instead of a sequential intermediate mechanism, SARM1 is unable to cleave cADPR in the absence of additive and in the presence of sodium citrate (Loring et al., 2020a) (Figure S2C–E). Interestingly, however, in the presence of PEG, SARM1 slowly cleaves cADPR, resulting in the time–dependent production of ADPR (Figure S2F–G).

**Figure 5.**
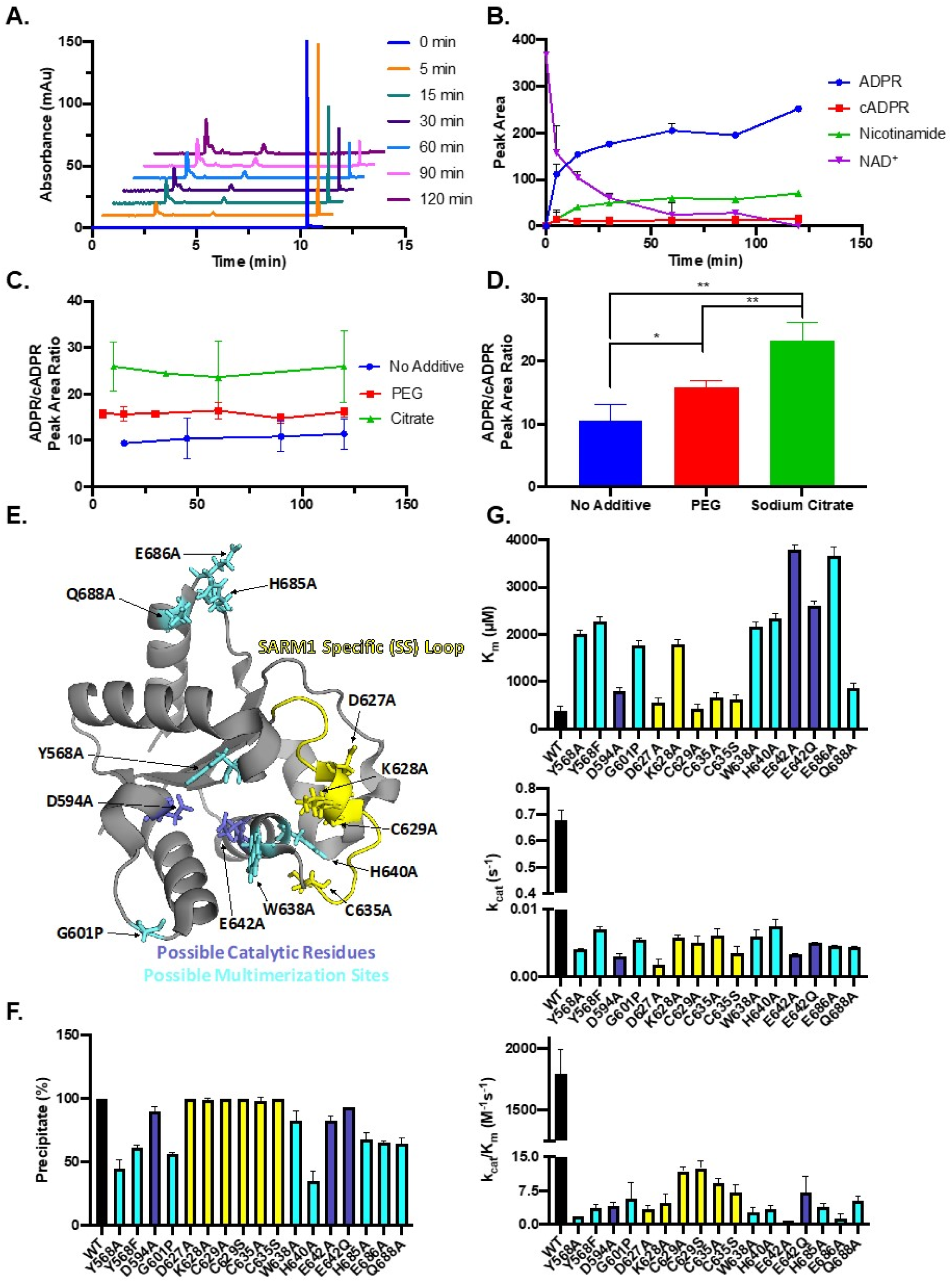
SARM1 Reaction Products and Site-directed Mutagenesis. (A) Representative time course of the SARM1-catalyzed reaction in the presence of PEG. The levels of ADPR, cADPR, nicotinamide, and NAD^+^ were detected by monitoring the absorbance at 254 nm. (B) Quantification of (A) (n =3). (C) ADPR/cADPR peak area ratios over time without additive, with PEG or with sodium citrate (n =3). (D) Average of ADPR/cADPR peak area ratios shown in panel C for no additive, PEG and sodium citrate SARM1 reactions, where * = P <0.0005 and ** = P<0.00005. (E) SARM1 mutants generated in this study. Highlighted residues include those present in the SARM1 Specific loop (yellow), potential catalytic residues (dark blue), and multimerization sites (light blue) (PDBID: 6o0q). (F) Precipitation of mutants compared to wild type SARM1 TIR domain (n =2). (G) *K*_m_, *k*_cat_, and *k*_cat_/*K*_m_ values for the mutants compared to wild type enzyme.

Next, we generated mutant constructs to understand how specific residues contribute to catalysis and multimerization. Residues were selected based on their proposed roles in multimerization, catalysis, or location in the SARM1 specific loop (Figure 5E). All the mutants examined showed a marked reduction in catalytic efficiency, *k*_cat_/*K*_m_, ranging from 100–2000–fold (Figure 5G). Interestingly, the most extreme effect was observed for the E642A mutant, which has previously been suggested to be a key catalytic residue based on structural alignments with other glycohydrolases (Essuman et al., 2017) (Figure 5G). Consistent with a role in multimerization the Y568, G601, H640, and H685 mutants showed reduced precipitation, suggesting that the loss in activity observed upon their mutation is likely due in part to a reduction in multimer formation (Figure 5F).

While *k*_cat_ values were reduced by ∼100–400–fold for all mutants examined, a subset showed a >4–fold increase in *K*_m_, including residues thought to be involved in multimerization, Y568, G601, W638, H640, and E686, and possible catalytic residues, K628 and E642. Several of these mutants have been previously characterized in the context of the purified protein and degenerating axons (Horsefield et al., 2019; Summers et al., 2016). Consistent with our results, they also found that TIR domain mutants E642A, Y568A, H685A, and BB loop mutations (D594A and G601P) reduce NAD^+^ hydrolase activity (Horsefield et al., 2019). Moreover, several of these mutations, including G601P, D627K, K628D, and C629S, were sufficient to prevent axonal degeneration post–axotomy, indicating that they render SARM1 non–functional (Summers et al., 2016)

Having performed a detailed study of SARM1 catalysis, we next sought to investigate how the liquid-to-solid phase transition enhances activity and why *k*_cat_, and therefore, catalytic efficiency increases with precipitant and enzyme concentration. To that end, we performed negative stain EM in the absence and presence of sodium citrate (500 mM) to determine how the precipitant affects activity and oligomeric state (Figure 6A–B). Surprisingly, we were able to detect the SARM1 TIR domain even in the absence of additive, which suggests the 20 kDa protein is forming a higher ordered oligomer. Notably, the diameter of the particles is on average ∼10 nm or ∼100 Å, which closely aligns with the diameter predicted for an octameric ring of the TIR domain. Supporting this finding are recent crystal and cryoEM structures of the SAM domains and full length protein, which demonstrate that SARM1 oligomerizes and forms an octameric ring (Bratkowski et al., 2020; Sporny et al., 2020; Sporny et al., 2019). In the presence of sodium citrate, SARM1 forms larger ring– like structures, potentially by wrapping around on itself (Figure 6B). These data suggest that precipitants like PEG 3350 and sodium citrate enhance SARM1 activity by inducing a liquid-to-solid phase transition, forming a higher ordered oligomer, and facilitating TIR domain multimerization.

**Figure 6.**
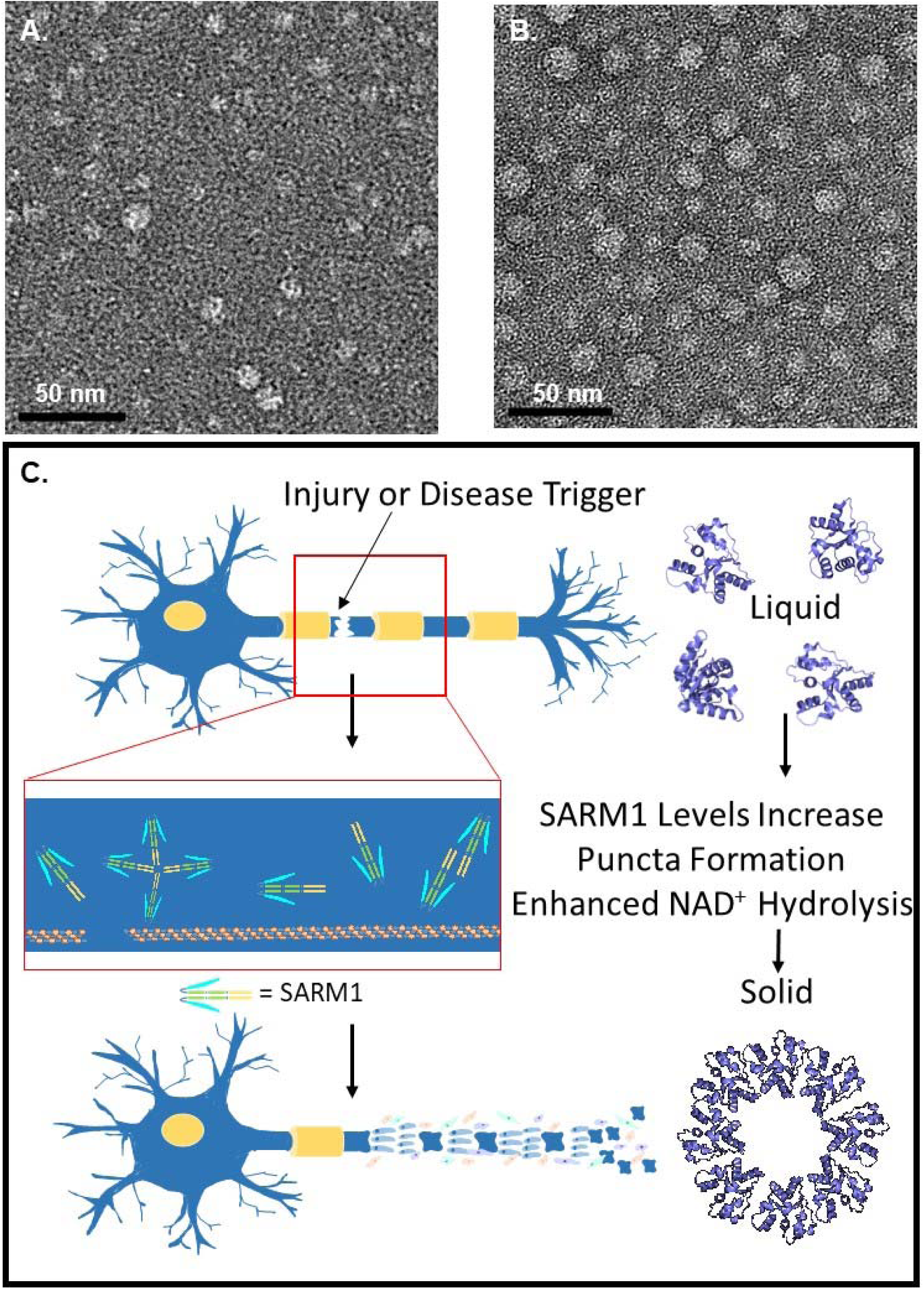
Negative stain EM images of SARM1 and Model for Enhancement of Activity. (A) Negative stain EM of SARM1 TIR domain. (B) Negative stain EM of SARM1 TIR domain with 500 mM sodium citrate. (C) Model for precipitant-mediated enhancement in SARM1 activity and subsequent axonal fragmentation.

## Conclusions

Together these experiments demonstrate that SARM1 activity is enhanced by the formation of an ordered oligomer and a liquid-to-solid phase transition. Furthermore, these data suggest that SARM1 activity is largely dependent on concentration and crowding alone and that as SARM1 multimerizes it becomes more active. SARM1 was previously thought to function as a dimer due to coimmunoprecipitation and FRET experiments; however, the SAM domains were recently crystallized as octamers (Gerdts et al., 2013; Horsefield et al., 2019; Summers et al., 2016) and other TIR domain-containing proteins (e.g., MAL) have been shown to form insoluble fibrils (Ve et al., 2017). Recent cryoEM structures of the full length enzyme in autoinhibited and active forms, suggests that SARM1 may actually form an octameric ring in both conformations, and that activity could be tied to a conformational change enabling multimerization of the TIR domains (Bratkowski et al., 2020; Sporny et al., 2020). The octameric ring formed by the SAM domains is reminiscent of that formed by inflammasomes and apoptosomes during an immune response or apoptosis. Our biochemical data supports the recent finding that SARM1 forms an octamer and that TIR domain multimerization is required to upregulate SARM1 activity. Our data further indicate that as SARM1 concentration increases, catalytic efficiency and turnover rate also increase. The fact that these changes correlate with the liquid to solid phase transition, suggests that as multimerization increases, SARM1 becomes a ‘better’ enzyme. In the context of a degenerating axon, this effect could translate into a feed–forward mechanism to induce degeneration. In fact, SARM1 protein levels have been documented to increase significantly in response to injury prior to Wallerian degeneration (Massoll et al., 2013), which is consistent with our data demonstrating that increasing SARM1 concentration enhances activity and with others showing upregulation of SARM1 is sufficient to induce axonal degeneration (Gerdts et al., 2013).

Based on these studies we propose a model whereby, increasing SARM1 concentration induces a phase transition that enhances SARM1 activity and is mimicked by PEG 3350 and sodium citrate. We hypothesize that the enhanced activity associated with this phase transition is due to the formation of an ordered oligomer like those recently observed in the crystal structure of the SAM domains and cryoEM structure of full length SARM1, and increased inter-TIR domain contacts (Figure 6C) (Bratkowski et al., 2020; Horsefield et al., 2019; Sporny et al., 2020). Interestingly, SARM1 forms higher ordered structures in neurons (both soma and dendrites) as punctate expression of this protein has been observed in numerous studies (Chen et al., 2011; Osterloh et al., 2012). Similar phase transitions have been observed in multiple signaling pathways. For example, the liquid phase condensation of cyclic GMP–AMP synthase (cGAS) activates innate immune signaling (Du and Chen, 2018) and liquid-to-solid phase transitions have been reported for FUS and huntingtin, during the formation of aggregates associated with ALS and Huntington’s disease (Patel et al., 2015; Peskett et al., 2018).

In summary, we show that SARM1 catalytic activity is enhanced by a liquid-to-solid phase transition reminiscent of that documented with other neurodegenerative disease–causing proteins. Therefore, inhibiting the liquid-to-solid transition could be a target for future therapeutic development for SARM1–associated diseases. In the context of a degenerating axon, we rationalize this phase transition as SARM1 exhibiting gradual NAD^+^ hydrolase activity upon initial activation until a critical threshold is reached whereby the activity increases exponentially, NAD^+^ is depleted, and the axon undergoes catastrophic fragmentation and granular disintegration. This could shed light on a built–in protective mechanism to control the toxic effects of SARM1; thus, ensuring the neuron has ample time to circumvent the degenerative path, but when committed to it, degeneration is executed in a positive feed–forward mechanism resulting in abrupt fragmentation and granular disintegration during the latter stages of Wallerian degeneration.

## Experimental Section

### Key Resources

Key resources, including constructs, primer sequences, chemicals, and antibodies are summarized in Table 3.

**Table S:**
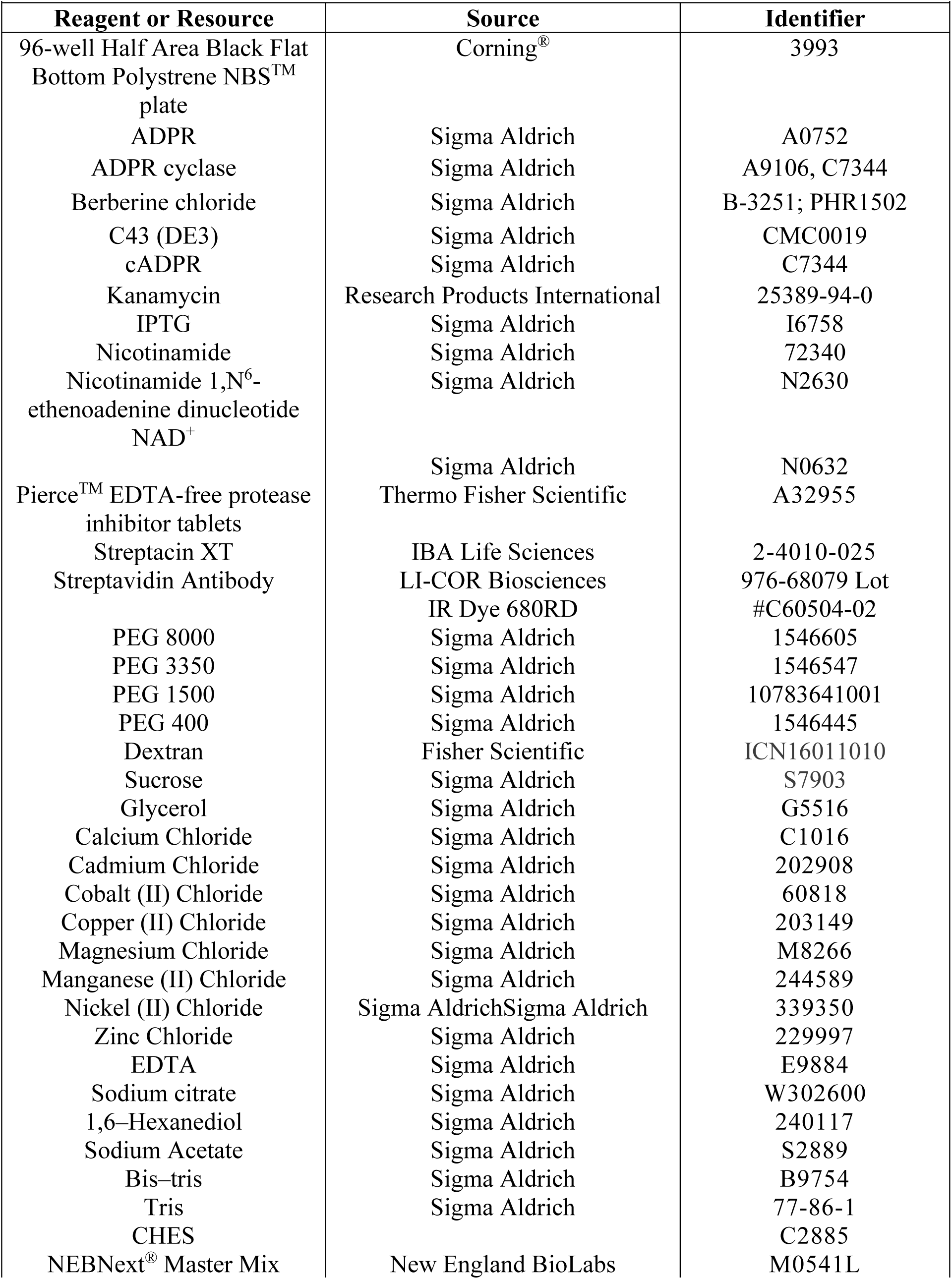
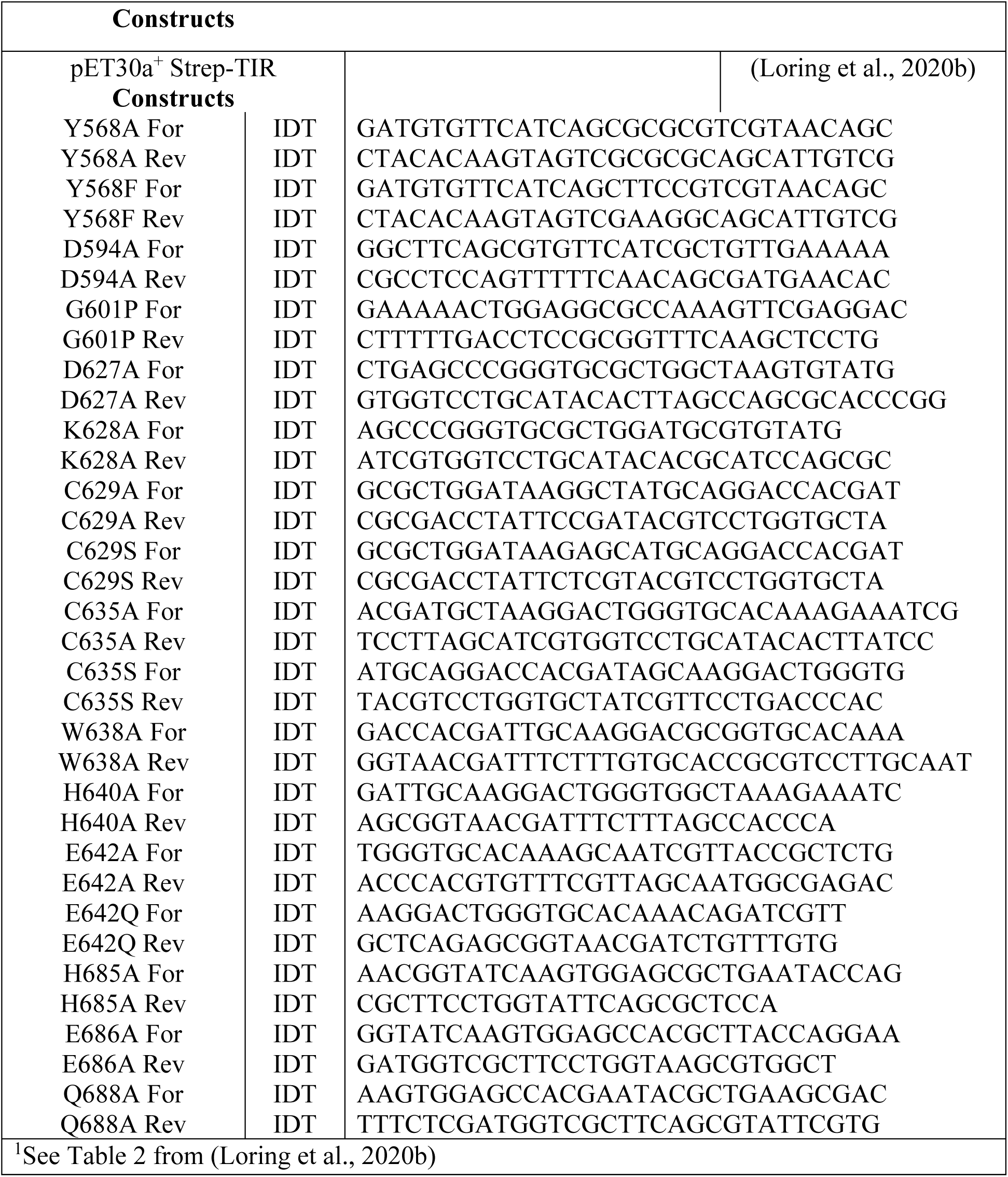
Key Resources.^1^.

### Recombinant SARM1 Expression and Purification

A recombinant human SARM1 TIR domain construct was bacterially expressed and purified as previously described (Loring et al., 2020a; Loring and Thompson, 2020b). Briefly, the TIR domain construct in the pET30a+ expression vector was transformed into chemically competent *Escherichia coli* C43 (DE3) cells and stored as a glycerol stock. Starter cultures were prepared by diluting glycerol stocks 1:1000 in LB media and grown at 37 °C overnight. The next day these starter cultures were diluted 400–fold in LB containing 50 µg/mL kanamycin and grown at 37 °C while shaking at 225 rpm until the culture reached an OD_600_ of 0.8. Protein expression was then induced by the addition IPTG (0.5 mM) and the incubation temperature was reduced to 16 °C for 16–18 h. Cells were then harvested by centrifugation at 3,000 x g for 15 min at 4 °C. The supernatant was removed and the pellet was resuspended in lysis buffer (100 mM HEPES pH 8.0, 200 mM NaCl, 10 % glycerol, 0.01% Tween 20) with Pierce™ EDTA–free protease inhibitor tablets (Thermo Scientific). The lysate was sonicated at an amplitude of 6 for 30 s (oscillating for 1 s on and 1 s off) for a series of 12 cycles using a Fisher Scientific Sonic Dismembrator sonicator (FB–705). After sonication, the lysate was clarified by centrifugation at 15,000 x g for 30 min at 4 °C. The clarified lysate was applied to pre–equilibrated Streptacin XT resin. The resin was then washed with Wash Buffer (50 mM HEPES pH 8.0 and 500 mM NaCl). SARM1 was eluted with Wash Buffer containing 50 mM biotin. The eluent was concentrated and injected on a HiLoad Superdec SUP75 (or SUP200) column for size exclusion chromatography using 50 mM HEPES pH 8.0 and 150 mM NaCl as the running buffer. Protein was concentrated using an Amicon 10 kDa cutoff centrifugal filter and protein concentration was determined using the Bradford assay.

### Determination of SARM1 Concentration in Lysates

The concentration of SARM1 in the lysates was determined by quantitative western blotting as previously described (Loring et al., 2020b). Briefly, serial dilutions of purified TIR domain protein (0 – 2 µM) were separated by SDS–PAGE along with lysate samples (1: 40 dilutions in duplicate). Next, proteins were transferred to nitrocellulose for western blotting. Protein was detected using a Streptavidin antibody (Table S3), which recognizes the N–terminal strep–tag. Band intensities were quantified (Licor) and used to generate standard curves, which were then applied to establish the concentration of SARM1 in the lysates.

### Fluorescent Assay

We applied a previously described continuous fluorescent assay to monitor SARM1 activity (Loring et al., 2020a). This assay employs an NAD^+^ analog, Nicotinamide 1, N^6^–ethenoadenine dinucleotide (ENAD), as a substrate. Upon hydrolysis and release of nicotinamide, etheno-ADPR (EADPR) fluoresces (λ_ex_= 330 nm, λ_em_= 405 nm). Enzymatic reactions were performed in Assay Buffer (20 mM HEPES pH 8.0 with 150 mM NaCl) and initiated by the addition of a 10x stock solution of ENAD in 96–well Corning^®^ Half Area Black Flat Bottom Polystrene NBS™ plates for a final reaction volume of 60 µL. EADPR fluorescence (λ_ex_= 330 nm, λ_em_= 405 nm) was detected in real time at 15 s intervals for the respective time period using a PerkinElmer EnVision 2104 Multilabel Reader in conjunction with Wallac EnVision Manager software. Activity was linear with respect to time and enzyme concentration under the conditions used. EADPR fluorescence (λ_ex_= 330 nm, λ_em_= 405 nm) was converted to molarity using an EADPR standard curve. Briefly, fixed concentrations of ENAD (0–400 µM) were treated excess ADP–ribosyl cyclase (Sigma Aldrich #A9106). The peak fluorescence intensities at each EADPR concentration were plotted against EADPR concentration to generate a standard curve.

Using this assay, the activity was monitored at each stage of the purification: crude lysate, clarified lysate, and purified protein, by adding SARM1 (300 nM) to Assay Buffer and then initiating the reaction with 100 µM ENAD. Activity was also measured by adding SARM1 (300 nM) back to empty pET30a+ vector C43 (DE3) lysate in Assay Buffer with 100 µM ENAD. These assays were performed for 15 min with readings taken every 15 s. The concentration dependence of purified SARM1 was also performed using this assay. The activity was monitored for serial dilutions of SARM1 (5 –35 µM) in Assay Buffer after the addition of 1 mM ENAD. Readings were taken at 15 s intervals for 30 min.

### Analytical Ultracentrifugation

Analytical ultracentrifugation sedimentation velocity analysis was performed at the UCONN Biophysics core. For these experiments, SARM1 TIR domain was studied at an OD_280_ of 0.3 and 1.0 in Assay Buffer at 20 °C and 50,000 rpm using absorbance optics with a Beckman-Coulter Optima analytical ultracentrifuge. Double sector cells equipped with quartz windows were used for analysis. The rotor was equilibrated under vacuum at 20 °C and after a period of ∼1 h at 20 °C, the rotor was accelerated to 50,000 rpm. Absorbance scans at 280 nm were acquired at 20 s intervals for ∼36 h. Data was analyzed with the Sedfit program using the direct boundary modeling program for individual data sets and model based numerical solutions to the Lamm equation. Continuous sedimentation coefficient c(s) distribution plots are sharpened, relative to other analysis methods, because the broadening effects of diffusion are removed by use of an average value for the frictional coefficient. The c(s) analyses were done at a resolution of 0.05 S, using maximum entropy regularization with a 95% confidence limit.

### Effect of Crowding Agents on Activity

Stock solutions of macroviscogens (PEG 8000, 3500, 1500, 400, Dextran) and microviscogens (Sucrose and Glycerol) were prepared at 50% w/v for the PEGs and 60% w/v for the remaining viscogens (Dextran, Sucrose, Glycerol). SARM1 was immersed in Assay Buffer at 10 µM without additive or with the addition of 30% w/v of each viscogen. After a 5 min incubation at RT, the reaction was initiated by the addition of 1 mM ENAD. The increase in fluorescence was monitored for 30 min as described above. The fluorescence was converted to EADPR produced using the EADPR standard curve. The slopes of these progress curves yielded the velocity of the reaction. The fold change relative to the no additive control was determined and the log fold change was plotted in GraphPad Prism. A representative progress curve is shown for SARM1 (20 µM) in Assay Buffer with and without 30% w/v PEG 3350. The reaction was initiated with 2 mM ENAD and readings were taken every 15 s for 15 min.

For the macroviscogens and glycerol, a dose dependence study was performed. SARM1 (10 µM) was immersed in Assay Buffer with 0, 10, 20, or 30% w/v of viscogen (PEG 8000, PEG 3350, PEG 1500, PEG 400, dextran, and glycerol). After a 5 min incubation at RT, ENAD was added (1 mM final) to initiate the reaction. The increase in fluorescence produced was monitored for 30 min and converted to velocity values at each viscogen concentration as described above. The fold change relative to the no additive control was determined and plotted in GraphPad Prism.

### Determination of Relative Viscosities

Viscosities were determined using a falling ball viscometer (Gilmont) and recording the time it takes for a steel ball to fall from the top line to the bottom line when the viscometer is filled with the respective solution. The viscosities were determined for water and 20% w/v solutions of dextran, sucrose, glycerol, and PEGs 8000, 3350, 1500, and 400. The relative viscosity was determined by calculating the fold–change relative to the viscosity of water.

### Effect of PEG 3350 on Enzyme Concentration Dependence

The enzyme concentration dependence studies were performed in Assay Buffer in the absence of PEG 3350 from 5–35 µM of SARM1 and in the presence of 12.5–25% w/v PEG 3350 with 300 nM–20 µM of SARM1. SARM1 was incubated under these conditions for 5 min at RT prior to initiation of the reaction by the addition of ENAD (1 mM). Fluorescence over time was recorded for 30 min and converted to velocity using a standard curve, as described above. Data was plotted in GraphPad Prism and fit to eq. 1,

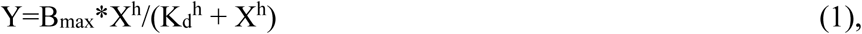

Where, X is the concentration, Y is specific binding, B_max_ is maximum binding, K_d_ binding coefficient, and h is the hill slope.

### Inhibitory Studies

To confirm that SARM1 behaves similarly in the presence and absence of viscogen, we assessed whether divalent metals affect SARM1 activity in the presence of 25% PEG 3350. Activity was recorded with and without 2 mM metals for purified SARM1 (2.5 µM) in Assay Buffer with 25% PEG 3350 after addition of 500 µM ENAD. The activity was also recorded for SARM1 (300 nM) in lysates with 2 mM metals after the addition of 50 µM ENAD. The activity relative to the no metal control was determined and values plotted in GraphPad Prism. For CdCl_2_ and CuCl_2_, IC_50_ values were determined at 500 nM, 1 µM, and 2.5 µM of SARM1 in Assay Buffer with 25% PEG 3350 and 500 µM ENAD.

We next determined IC_50_ values for zinc chloride and berberine chloride in the presence and absence of 25 % w/v PEG3350. For the experiments with PEG, SARM1 (2.5 µM) was combined with 25% PEG 3350 in Assay Buffer. SARM1 (20 µM) was studied in Assay Buffer for the experiments without PEG 3350. Inhibitor (zinc chloride or berberine chloride) was added at concentrations from 0–500 µM and incubated for 5 min at RT prior to initiating the reaction with ENAD (100 µM final). The data were converted to velocity as described above and then normalized to the activity without the inhibitor. IC_50_ values were calculated by fitting the normalized inhibition data to eq. 2,

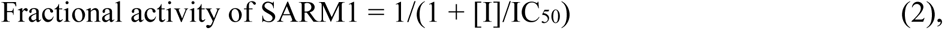

in GraphPad Prism. [I] is the concentration of inhibitor, IC_50_ is the concentration of the inhibitor at half the maximum enzymatic activity, and fractional activity of SARM1 is the percent activity at the respective inhibitor concentration.

To further assess whether the results in lysate are recapitulated with pure protein in the presence of viscogen, mechanism of inhibition studies were repeated with zinc chloride and berberine chloride. For these experiments, SARM1 (2.5 µM) was combined with 25% w/v PEG 3350 in Assay Buffer with either zinc chloride (0–2.5 µM) or berberine chloride (0–200 µM). Reactions were initiated with ENAD (0–2 mM). Reactions were monitored for 30 min as described above. The initial velocities were compiled to produce Michaelis Menten curves at each inhibitor concentration and fit to equations for competitive (eq 3), noncompetitive (eq 4), and uncompetitive inhibition (eq 5),

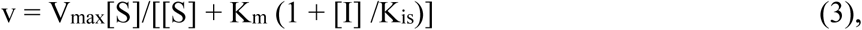

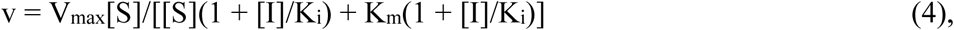

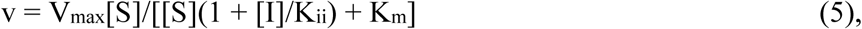

in GraphPad Prism. *K*_ii_ is the intercept *K*_i_ and *K*_is_ is the slope *K*_i_. The best fits were determined on the basis of a combination of visual and quantitative analysis.

### PEG 3350 and Sodium Citrate Precipitate SARM1

SARM1 (20 µM) was incubated at room temperature for 10 min in Assay Buffer with or without 25% PEG 3350. At which time, the solutions were separated by centrifugation at 21,000 x g for 10 min at 4 °C. The supernatant was removed and pellet resuspended in Assay Buffer plus 25% PEG 3350. The pre-centrifugation, supernatant, and resuspended pellet samples were assayed by the addition of 1 mM ENAD and monitored for 30 min as described above. Coomassie gels were run on aliquots of each of the fractions. The same experiment was also performed at 5, 10, and 20 µM SARM1 in the presence of 0, 10, 17.5, and 25% PEG 3350 in assay buffer. Briefly, SARM1 (5, 10, and 20 µM) was incubated in Assay Buffer containing (0, 10, 17.5, or 25%) PEG 3350 for 10 min at RT. Then the solutions were separated by centrifugation at 21,000 x g for 10 min at 4 °C. The supernatant was removed, and pellet resuspended in Assay Buffer plus the respective PEG concentration. Each fraction was assayed for activity via the addition of 1 mM ENAD and presence of SARM1 confirmed by commassie gel.

These centrifugation experiments were also performed with sodium citrate at concentrations of 100, 250, 500, 750, and 1000 mM in Assay Buffer. Specifically, SARM1 (5 µM) was incubated at room temperature for 10 min in Assay Buffer with different concentrations of sodium citrate. After which, the samples were separated by centrifugation at 21,000 x g for 10 min at 4 °C. The supernatant was removed, and the pellet resuspended in Assay Buffer plus sodium citrate. The resuspended pellet samples were assayed via the addition of 1 mM ENAD and monitored for 30 min as described above. The pellet and supernatant samples were run on a coomassie gel for each sodium citrate concentration.

### Effect of 1,6–Hexanediol on Activity

The effect of 1,6–hexanediol, an aliphatic alcohol, on SARM1 activity was assessed. SARM1 (20 µM) was added to Assay Buffer containing 0–2% 1,6–hexanediol and the reaction was initiated via the addition of 1 mM ENAD. The reaction was monitored for 1 h as described above. The effect of 1,6–hexanediol was also assessed in the presence of precipitants. SARM1 (2.5 µM) was added to Assay Buffer with 25% PEG 3350 or 500 mM sodium citrate with and without 2% 1,6–hexanediol. The reaction was initiated via the addition of 1 mM ENAD and monitored for 30 min as described above.

### Determination if Precipitation is Reversible

To investigate whether SARM1 precipitation is reversible, SARM1 (10 µM) was added to Assay Buffer with either 25% PEG 3350 or 500 mM sodium citrate. Samples were then separated by centrifugation at 21,000 x *g* for 10 min at 4 °C. The supernatant was removed, and the pellet was resuspended in buffer alone or buffer plus additive. The activity of the different fractions were assessed at 5 µM SARM1 final and 500 µM ENAD as described above. To confirm that there was not a precipitate present after resuspension in buffer alone but still present after resuspension with buffer plus additive, samples were again separated by centrifugation at 21,000 x *g* for 10 min at 4 °C and visually inspected.

### Effect of pH on SARM1 Activity

To determine if pH affects the activity of pure enzyme, buffer stocks (4x) were prepared at 80 mM and 600 mM NaCl. The following buffers were used: Sodium Acetate pH 4, 4.5, 5, and 5.5; Bis–tris pH 6.0 and 6.5; Tris pH 7.0, 7.5, 8.0 and 8.5; and CHES pH 9.0. For the assays, the 4x buffer stocks were diluted with water and 25% PEG 3350 for a final concentration of 20 mM and 150 mM NaCl and the pH was confirmed after dilution. SARM1 (2.5 µM) was added to the buffer plus PEG solution and incubated at RT for 5 min prior to initiation of the reaction with ENAD (0–2 mM). These experiments were performed three times in duplicate. The fluorescence was recorded as described above for 1 h. The initial velocities were compiled in GraphPad Prism and data for each pH was fit to the Michaelis Menten equation (eq. 6),

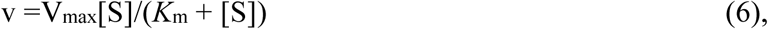

V_max_ is the maximum velocity, [S] is the concentration of substrate, and *K*_m_ is the substrate concentration at half the maximum velocity. The log *K*_m_, *k*_cat_ and *k*_cat_/*K*_m_ values were plotted against pH and fit to a bell shaped equation (eq. 7),

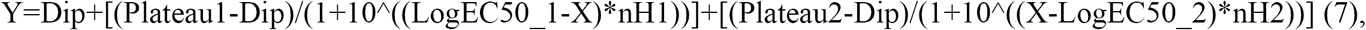

X is the Log pH, Y is the response, Plateau 1 and 2 are the initial and final plateaus, Dip is the plateau level between phases, Log EC50_1 and 2 are equivalent to pka1 and 2, and nH1 and 2 are the slope factors.

Percent precipitation was also assessed from pH 4–9 for SARM1 with and without PEG 3350. SARM1 (2.5 µM) in 1x buffer alone or with 25% PEG 3350 was incubated at RT for 10 min. At which time, the samples were separated by centrifugation at 21,000 x g for 10 min at 4 °C. The supernatant was removed, and the pellet resuspended in buffer alone or with 25% PEG 3350. Supernatant and pellet samples with and without PEG 3350 were run on an SDS–PAGE gel. Stain free images were obtained, and bands were quantified using ImageJ software. The percent precipitation values were averaged for the experiment performed in duplicate and the values fit to eq. 7.

### Effect of PEG 3350 on Steady State Kinetics

The steady state kinetic reactions were performed in Assay Buffer with varied PEG 3350 (0–25%) at a constant SARM1 concentration (10 µM). Reactions were incubated at RT for 5 min and initiated by the addition of ENAD (0–2 mM final). Fluorescence was recorded as described above for 1 h. The initial velocities were compiled and fit to eq. 6 in GraphPad Prism to produce Michaelis Menten curves at each PEG 3350 concentration.

The fold change was calculated for the *K*_m_, *k*_cat_, and *k*_cat_/*K*_m_ values at 10 µM SARM1 with the addition of 25% w/v PEG 3350 compared to the values without PEG 3350 and the log values plotted using GraphPad Prism. The *K*_m_, *k*_cat_ and *k*_cat_/*K*_m_ values were also plotted against PEG 3350 concentration and trend evident with a spline (*K*_m_) or sigmoidal fit (*k*_cat_ and *k*_cat_/*K*_m_) (eq. 8),

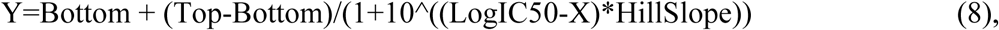

X is the log(concentration), Y is the response, Top and Bottom are plateaus, log IC_50_ is the log of the concentration of the inhibitor at half the maximum enzymatic activity.

These experiments were also performed at constant PEG 3350 (25%) and varied SARM1 concentration (500 nM, 1 µM, 2.5 µM, 5 µM and 10 µM), and without PEG 3350 at 10 and 20 µM SARM1. SARM1 was incubated in Assay Buffer with or without PEG 3350 at RT for 5 min and then the reaction was initiated by the addition of ENAD (0–2 mM final). Fluorescence was recorded as described above for 1h. The initial velocities were compiled to produce Michaelis Menten curves at each SARM1 concentration with and without PEG 3350. These slopes were plotted in GraphPad Prism and fit to the Michaelis Menten equation (eq. 6). From this analysis, the *K*_m_, *k*_cat_ and *k*_cat_/*K*_m_ values were plotted against enzyme concentration and the trend is shown with a spline fit.

### LC–MS Based Assay

ADPR, cADPR, nicotinamide, and NAD^+^ standards were prepared at 100 µM in Assay Buffer. Purified SARM1 (20 µM) was treated with 100 µM NAD^+^ in Assay Buffer and the reaction was monitored over time. Samples were injected at 0, 15, 40, 90, and 120 min onto an InfinityLab Poroshell 120 EC-18 column (4.6 x 50 mm, 2.7 mircron, Agilent #699975–902T) at a flow rate of 0.4 mL/min. Metabolites were eluted with a gradient of 100% H_2_O with 0.01% formic acid from 0 to 6 min, 90% H_2_O with 0.01% formic acid and 10% acetonitrile with 0.1% formic acid from 12 to 14 min, and then 100% H_2_O with 0.01% formic acid from 14 to 20 min. The metabolites were detected with a quadrupole mass spectrometer (Agilent, G6120B single quad, ESI source, 1260 Infinity HPLC) in the positive ion mode. The absorbance at 254 nm was plotted over time at each reaction time point in GraphPad Prism. Reaction time courses were also performed as described above, in the presence of 25% PEG 3350 or 500 mM sodium citrate at 2.5 µM SARM1. The reactions were injected at time 0, 5, 15, 30, 60, 90, and 120 min for PEG 3350, and 0, 10, 30, 60, 90, 120, and 150 min for sodium citrate. Peak areas were quantified at an absorbance of 254 nm and their accumulation or consumption plotted over time. The ratio of ADPR to cADPR was also quantified for each condition over time and averaged for the entire time course. A student’s t–test was performed to determine if the differences in ADPR to cADPR ratios in the absence or presence of additive were statistically significant (P value = 0.00036 for No PEG 3350/PEG 3350, 0.0000024 No sodium citrate/sodium citrate, and 0.000014 for PEG 3350/sodium citrate.

Reaction time courses were also performed with 100 µM cADPR and SARM1 alone (20µM), and with 25% PEG 3350 or 500 mM sodium citrate at 5 µM enzyme in Assay Buffer. For SARM1 alone and SARM1 with sodium citrate, the reaction was monitored at time 0, 2 h and 6 h. SARM1 with PEG 3350 was recorded at time 0, 10, 30, 60, 90, and 150 min. Samples at each time point were injected onto an InfinityLab Poroshell 120 EC-18 column (4.6 x 50 mm, 2.7 mircron, Agilent #699975–902T) at a flow rate of 0.4 mL/min. Metabolites were eluted with a gradient of 100% H_2_O with 0.01% formic acid from 0 to 6 min, 90% H_2_O with 0.01% formic acid and 10% acetonitrile with 0.1% formic acid from 12 to 14 min, and then 100% H_2_O with 0.01% formic acid from 14 to 20 min. The metabolites were detected with a quadrupole mass spectrometer (Agilent, G6120B single quad, ESI source, 1260 Infinity HPLC) in the positive ion mode. The absorbance at 254 nm was plotted over time at each reaction time point in GraphPad Prism. The peak areas were quantified for ADPR and cADPR at an absorbance of 254 nm and the average was over time was plotted in GraphPad Prism.

### TIR Domain Mutants

TIR domain mutants were generated using the pET30a+ TIR domain construct as a template (Table 3). To mutagenize the plasmid by PCR, template DNA (2 ng/µL) was combined with forward and reverse primers (0.5 µM final, Table 3) in 1xNEBNext^®^ Mastermix, which contains the Q5 high fidelity polymerase, deoxynucleotides, and 2 mM MgCl_2_.The PCR protocol was as follows: (1) initial denaturation at 98 °C for 1 min, (2) denaturation at 98 °C for 30 s, (3) annealing at 54 – 56 °C for 45 s, (4) extension at 68 °C for 1 min and (5) final extension at 68 °C for 5 min. Steps 2 – 4 were repeated 30 times. Site–directed mutagenesis was confirmed by Sanger sequencing (Genewiz).

Precipitation of mutants was assessed by combining SARM1 (10 µM) with 25% PEG 3350 in Assay Buffer. Samples were separated by centrifugation at 21,000 x g for 10 min at 4 °C. The supernatant was removed, and the pellet was resuspended in buffer. Coomassie gels were run of the supernatant and pellet fractions in duplicate and band intensities were quantified with ImageJ and percent precipitation plotted in GraphPad Prism.

Steady state kinetic parameters were determined at 5 µM mutant SARM1 and 25% PEG 3350 in Assay Buffer. Reactions were initiated by the addition of ENAD (0–2 mM). Fluorescence was recorded as described above for 1 h. The initial velocities were compiled in GraphPad Prism and fit to the Michaelis Menten equation (eq. 6). From this analysis, the *K*_m_, *k*_cat_ and *k*_cat_/*K*_m_ values were obtained for each mutant and plotted in GraphPad Prism.

### Negative Stain Electron Microscopy on SARM1

We performed negative stain EM on SARM1 TIR domain (10 µg/mL) in Assay Buffer with and without sodium citrate (500 mM). Samples were processed by the core facility at UMass Medical School. Samples were fixed with uranyl acetate and imaged on the Philips CM120 microscope.

## AUTHOR INFORMATION

### Corresponding Author

Mailing address: Department of Biochemistry and Molecular Pharmacology, University of Massachusetts Medical School, LRB 826, 364 Plantation Street, Worcester MA 01605. Tel.: 508-856-8492. Fax: 508-856-6215.

### Notes

The authors declare no competing financial interest(s),

### Funding Sources

This work was supported in part by NIH grants R35 GM118112 (P.R.T.) and F31NS108610 (H.S.L.).

## Supporting Information

**Figure S1.**
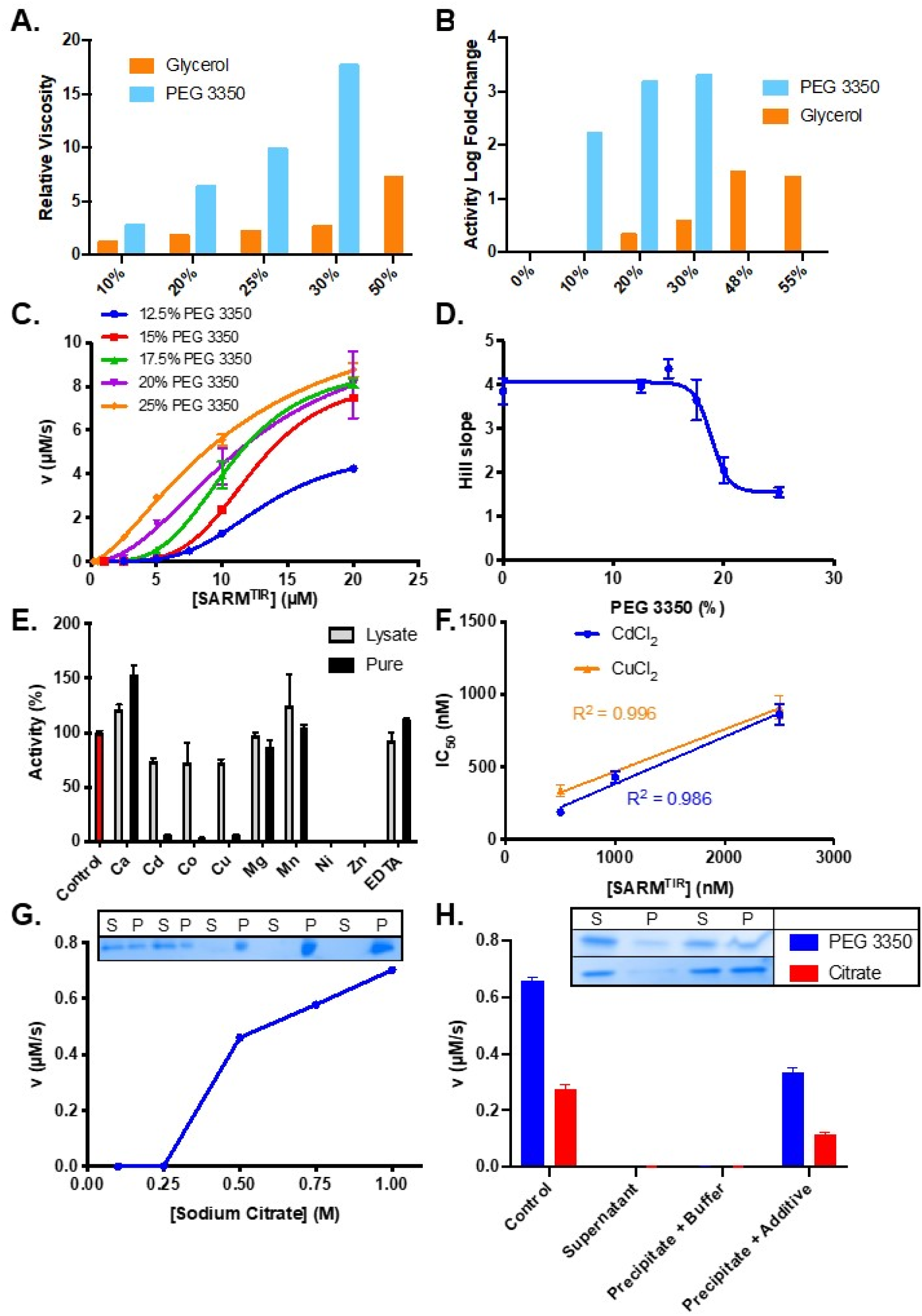
Effect of Additives on Viscosity and Activity. (A) Viscosities of glycerol and PEG 3350 at 10–50% w/v. (B) Fold increase in activity 0–55% glycerol. (C) Activity at 0.25 – 20 µM SARM1 with 12.5 – 25% PEG 3350 fit to equation 1. (D) Fitting to equation 1 gives hill slopes for each concentration of PEG. (E) Inhibition of SARM1 (lysate, 300 nM and pure, 2.5 µM with 25% PEG 3350) by divalent metals (2 mM). (F) IC_50_ of CuCl_2_ and CdCl_2_ with SARM1 at three concentrations of enzyme. (G) Activity of SARM1 at 100, 250, 500, 750, and 1000 mM sodium citrate, with coomassie gel depicting SARM1 in supernatant (S) and pellet (P) fractions. (H) Activity of pre-centrifugation control, supernatant, precipitate resuspended in buffer and precipitate resuspended in buffer plus additive. Coomassie gels are shown for each fraction with PEG 3350 and sodium citrate. Full gel provided in Figure S3E.

**Figure S2.**
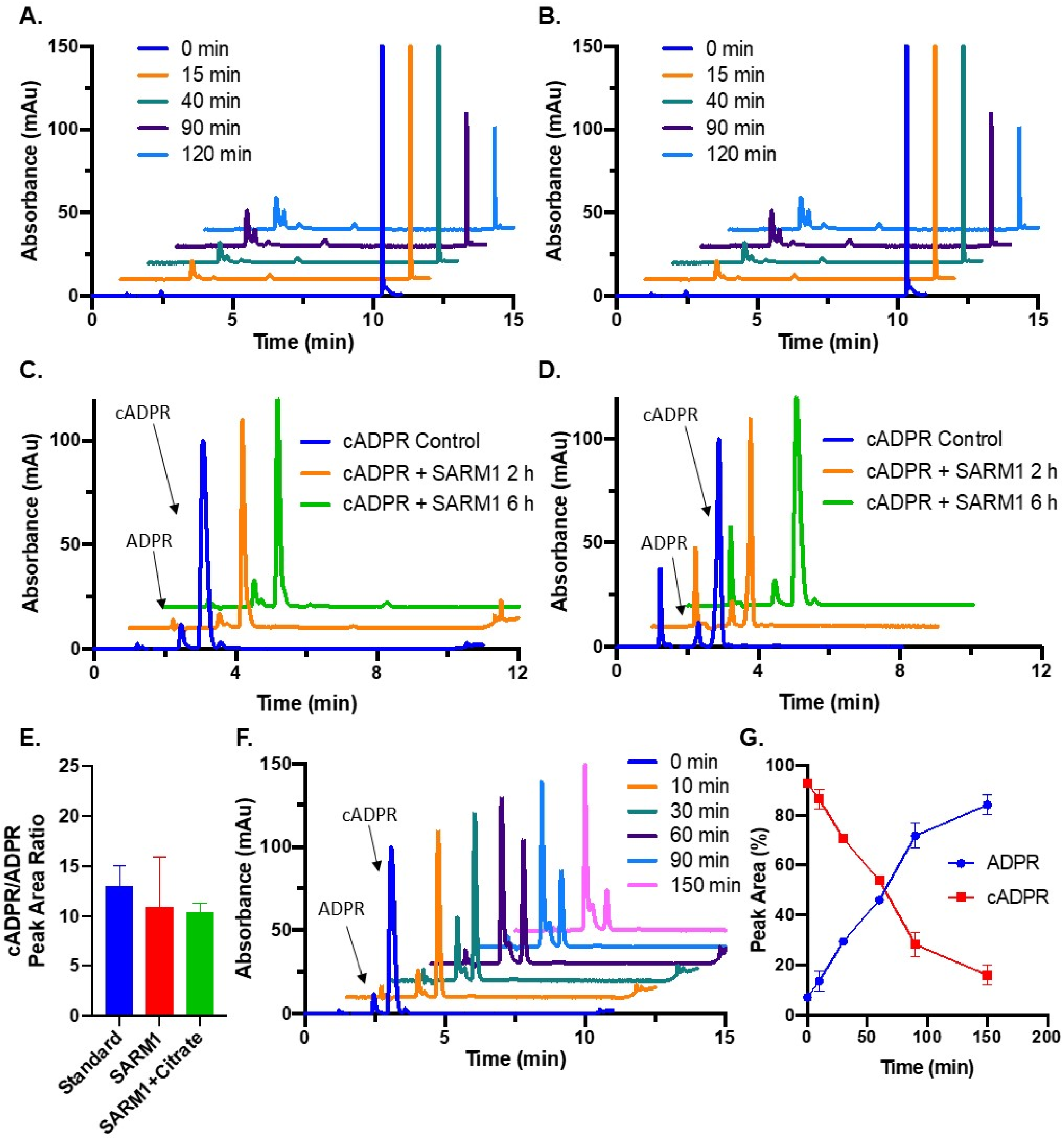
LC–MS Assays. (A) Representative SARM1 reaction time course treated with 100 µM NAD^+^ showing the absorbance at 254 nm for NAD^+^, nicotinamide, ADPR, and cADPR over time without additive. (B) Representative SARM1 reaction time course with sodium citrate treated with 100 µM NAD^+^ showing the absorbance at 254 nm for NAD^+^, nicotinamide, ADPR, and cADPR over time. (C) Absorbance at 254 nm for SARM1 (20 µM) treated with 100 µM cADPR without additive for 2 h and 6 h, shown with 100 µM cADPR standard. (D) Absorbance at 254 nm for SARM1 (20 µM) treated with 100 µM cADPR with 500 mM sodium citrate for 2 h and 6 h, shown with 100 µM cADPR standard. Quantification of cADPR/ADPR peak area ratios in control, SARM1 reaction, and SARM1 reaction with sodium citrate. Experiments were performed in duplicate. (E) Representative SARM1 reaction time course with PEG treated with 100 µM cADPR showing the absorbance at 254 nm for ADPR and cADPR over time. (F) Quantification of ADPR and cADPR peak areas at an absorbance of 254 nm. Experiments were performed in duplicate.

**Figure S3.**
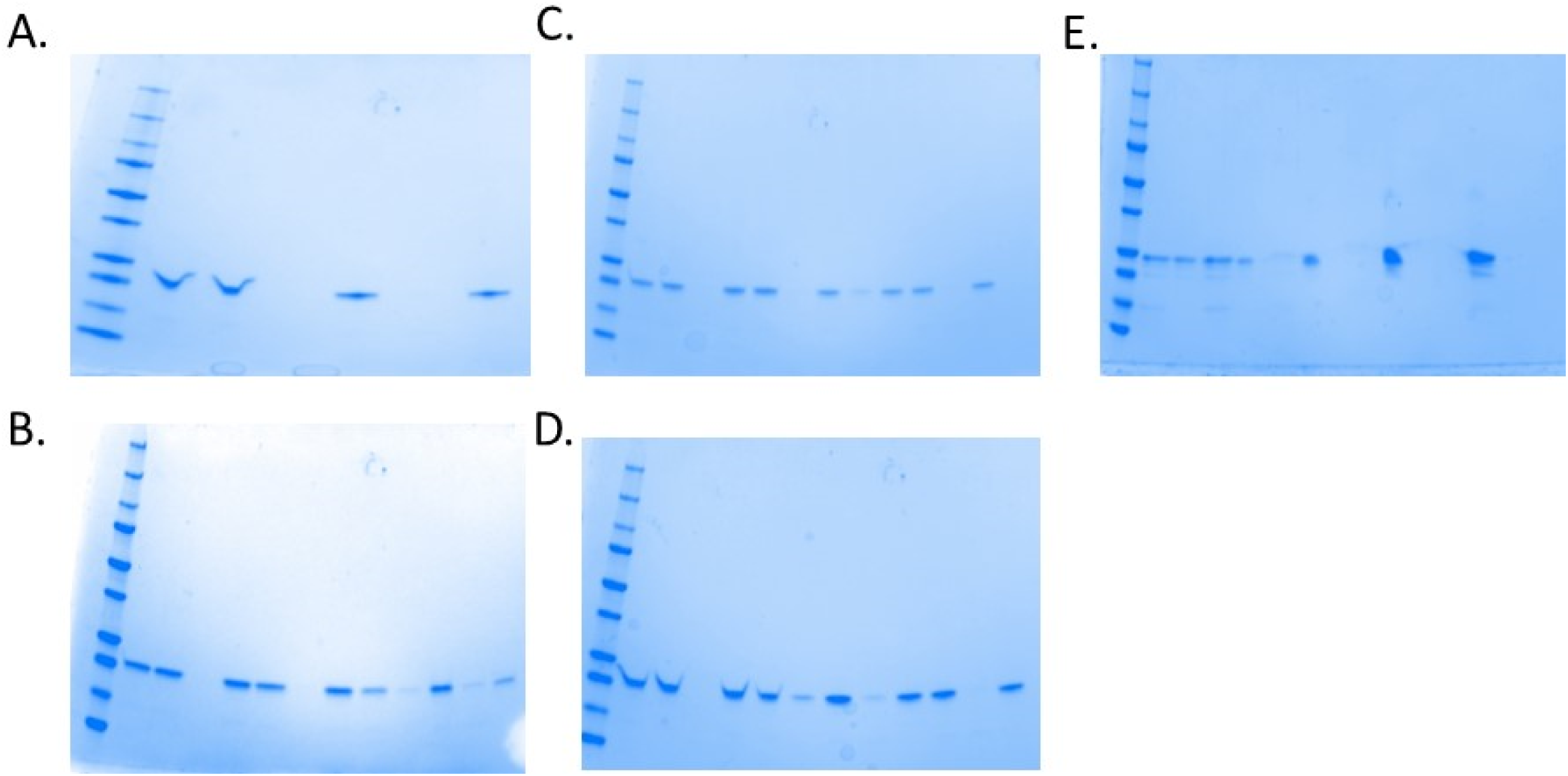
Full gels of those shown in Figure 3 and Figure S1. (A) Pre-centrifugation, supernatant, and pellet fractions of 20 µM SARM1 without and with 25% PEG. (B) Pre-centrifugation, supernatant, and pellet fractions at 5 µM SARM1 and 0, 10, 17.5, and 25% PEG 3350. (C) Pre-centrifugation, supernatant, and pellet fractions at 10 µM SARM1 and 0, 10, 17.5, and 25% PEG 3350. (D) Pre-centrifugation, supernatant, and pellet fractions at 20 µM SARM1 and 0, 10, 17.5, and 25% PEG 3350. (E) Full gel of image of the gel in Figure S1E, showing SARM1 in supernatant and pellet fractions with 0.1-1M sodium citrate.

**Table S1.**
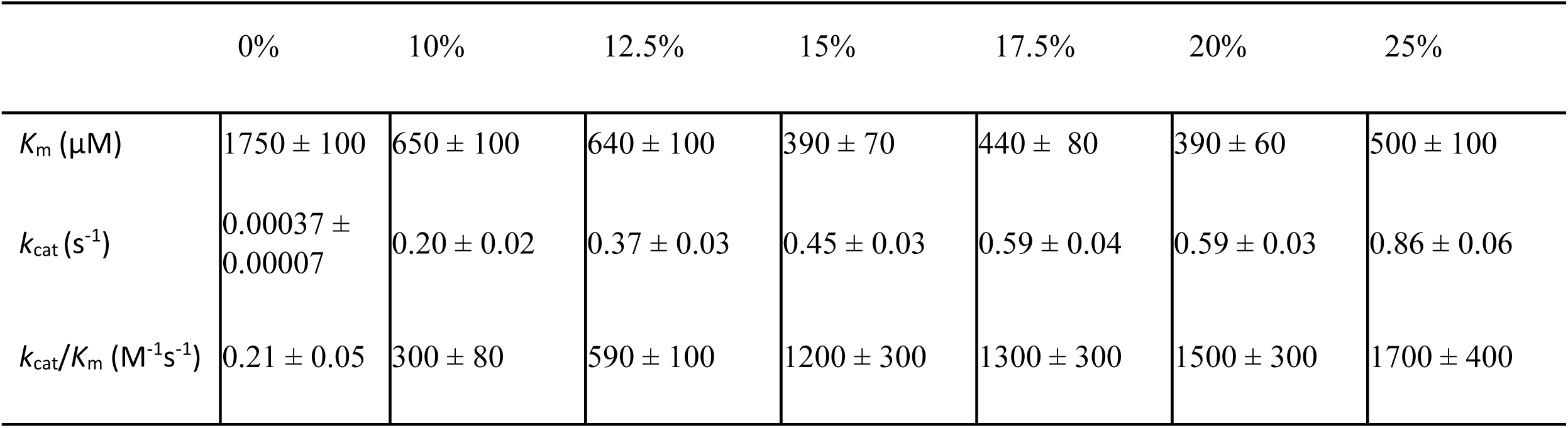
The Steady-State Kinetic Parameters for Pure Wild Type SARM1 TIR (10 µM) with Varied PEG.

|  | 0% | 10% | 12.5% | 15% | 17.5% | 20% | 25% |
| --- | --- | --- | --- | --- | --- | --- | --- |
| $K_m$ (μM) | 1750 ± 100 | 650 ± 100 | 640 ± 100 | 390 ± 70 | 440 ± 80 | 390 ± 60 | 500 ± 100 |
| $k_{cat}$ ($s^{-1}$) | 0.00037 ± 0.00007 | 0.20 ± 0.02 | 0.37 ± 0.03 | 0.45 ± 0.03 | 0.59 ± 0.04 | 0.59 ± 0.03 | 0.86 ± 0.06 |
| $k_{cat}/K_m$ ($M^{-1}s^{-1}$) | 0.21 ± 0.05 | 300 ± 80 | 590 ± 100 | 1200 ± 300 | 1300 ± 300 | 1500 ± 300 | 1700 ± 400 |

**Table S2.** The Steady-State Kinetic Parameters for Pure Wild Type SARM1 TIR with Varied Enzyme Concentration.

| | 500 nM | 1 μM | 2.5 μM | 5 μM | 10 μM | 10 μM | 20 μM |
| --- | --- | --- | --- | --- | --- | --- | --- |
|  | PEG | PEG | PEG | PEG | PEG |  |  |
| $K_m$ (μM) | $1600 \pm 600$ | $800 \pm 30$ | $230 \pm 30$ | $380 \pm 70$ | $500 \pm 90$ | $1700 \pm 600$ | $1200 \pm 200$ |
| $k_{cat}$ ($s^{-1}$) | $0.24 \pm 0.04$ | $0.19 \pm 0.03$ | $0.51 \pm 0.01$ | $0.68 \pm 0.04$ | $0.86 \pm 0.06$ | $0.0004 \pm 0.0001$ | $0.0023 \pm 0.0002$ |
| $k_{cat}/K_m$ ($M^{-1}s^{-1}$) | $150 \pm 70$ | $240 \pm 10$ | $2200 \pm 30$ | $1800 \pm 70$ | $1700 \pm 70$ | $0.24 \pm 0.01$ | $1.9 \pm 1$ |

## Notes

### Competing Interest Statement

The authors have declared no competing interest.

